# An information theoretic approach to evaluating social behaviour in mice using automated home-cage monitoring

**DOI:** 10.64898/2026.11.23.767392

**Authors:** Tiffany Chien, Darren J. Fernandes, John G. Sled, Jason P. Lerch

## Abstract

**Background:** Social behaviour in mice is commonly assessed using standardized assays such as the three-chambered task, which provide controlled measures of sociability but often capture behaviour over short timeframes that may not capture the complex behavioural repertoire exhibited in the home environment. Although automated home-cage monitoring enables continuous, longitudinal assessment of group-housed mice with minimal human interference, methods for quantifying social behaviour in this complex setting remain limited. Information theory offers a framework for measuring statistical dependence between animals but requires one to account for dependence related to the cage environment that arise even in the absence of social interaction.

**Methods:** We developed an information-theoretic approach to quantify social dependence using longitudinal RFID-based positional data from group-housed mice monitored with automated home-cage monitoring system. Mutual information (MI) between cagemates was evaluated against a constrained null model that disrupted within-cage dependence while preserving individual movement patterns. We used two strains with known differences in sociability (C57BL/6J and BTBR) to assess sensitivity to various temporal sampling parameters, test-retest reliability, and correspondence with manually video-scored social behaviours. We further examined variation across genetic backgrounds and investigated relationships between home-cage measures, conventional behavioural assays (three-chamber sociability and open-field behaviour), and brain structure using *in vivo* MRI.

**Results:** Observed MI was distinguishable from our null distribution, validating its use to quantify statistical dependence beyond that expected from shared cage constraints. MI recapitulated known differences in sociability between C57BL/6J and BTBR mice, showed high test-retest reliability, and correlated with independently video-scored social behaviours. Home-cage social measures also varied across genetic backgrounds and captured strain differences separate from three-chamber sociability. Furthermore, *in vivo* MRI revealed distinct neuroanatomical patterns for home cage compared to three-chamber sociability measures, including brain regions implicated in social behaviour. Comparisons with open-field measures further indicated that home-cage information-theoretic measures capture behavioural variation not fully represented by conventional measures of locomotion and exploration. Finally, social coordination exhibited distinct temporal dynamics across the circadian cycle.

**Conclusion:** This study establishes and validates an information-theoretic framework for quantifying social dependence from longitudinal home-cage positional data. By accounting for dependence arising from shared environmental constraints and demonstrating reproducibility and correspondence with observed social behaviour, this approach provides a complementary method for characterizing social behaviour in mice. Its application across genetic backgrounds and temporal contexts offers opportunities to investigate behavioural variation beyond the restricted sampling of conventional assays.

## INTRODUCTION

Social interactions are fundamental to mammalian behaviour, enabling communication, relationship formation, and navigation of complex social environments. Impairments in social behaviour are characteristic of several neuropsychiatric and neurodevelopmental disorders, including autism spectrum disorders (ASD), attention deficit hyperactivity disorder (ADHD), obsessive compulsive disorders (OCD), and schizophrenia.^1,2^ Rodents provide valuable models for investigating the neural mechanisms underlying these conditions because they exhibit similar complex social behaviors including empathy, recognition, affiliation, and hierarchy formation,^3^ and can be easily modified to exhibit core behavioural features of these disorders.

While social behavior is inherently dynamic and reciprocal, rodent social behaviour phenotyping has traditionally relied on short-duration, experimenter-administered tests conducted in unfamiliar environments, often during the inactive phase of the circadian cycle.^4^ The three-chamber task, for example, is widely used to assess sociability by measuring an animal’s preference for a novel conspecific over a non-social stimulus.^5,6^ Although readily automated and suitable for high-throughput testing, its design inherently restricts reciprocal physical interaction.^7,8^ Further, it is often difficult to determine whether observed behaviours represent genuine affiliative social motivation or are instead driven by exploration, novelty seeking, or territorial investigation.^9,10^ Reciprocal social interaction assays allows more direct and complex interactions, including sniffing, allogrooming, chasing, following,^11^ while partition and partner preference tests evaluate social affiliation, recognition, and memory through measures of proximity and preference for social stimuli.^11^ However, these assays remain constrained by brief testing periods, repeated handling, novel environmental contexts, and potential aggression or dominance effects.^11^ More broadly, conventional assays sample behaviour within restricted temporal windows and often reduce complex social interactions to a limited set of predefined behavioural measures, which may be particularly restrictive for studying social behaviours that emerge through quick, reciprocal exchanges between multiple individuals over time. Furthermore, differences in experimental design and analysis have contributed to limited reproducibility across studies and laboratories.^4,12–14^

Automated home cage monitoring (HCM) addresses these temporal and contextual limitations by enabling continuous observation of animals within their familiar housing environment.^15^ These systems offer a minimally invasive, high-throughput, and scalable platform for assessing spontaneous behaviour over extended periods across both the light and dark cycles, providing longitudinal measurements that are difficult to obtain using conventional behavioural assays.^16^ For example, home-cage monitoring has been used to characterize circadian activity patterns, estrous-related behavioural changes, and individual differences in spontaneous behaviour, and may provide improved opportunities for monitoring behavioural responses to interventions over time.^17–19^ Individual identification further enables persistent behavioral traits and social dynamics to be captured that may be otherwise missed by testing within a single session, such as characterization of individual behavioural phenotypes, pairwise interactions, and collective social organization.^15^ Several studies have also demonstrated the advantages of using automated HCM over traditional behavioral tests for improving reproducibility across labs.^18,20,21^ Together, these approaches highlight the potential of longitudinal home-cage monitoring to capture social dynamics over time.

An additional challenge involves the quantification of social behaviour from continuous, multi-animal recordings. Characterizing social behaviour has traditionally relied on automated classification or predefined behaviors such as proximity, locomotion, chasing, or avoidance.^22,23^ These approaches provide useful and interpretable descriptions of specific behaviours, but may not fully capture the dynamic relationships that emerge spontaneously between multiple animals.^15,17,24^ This is particularly challenging when several animals interact simultaneously, and behavioural states change rapidly and overlap, while occlusion from bedding, food, enrichment, and other objects can further complicate video-based analysis in the home cage. Instead of scoring social interaction according to a predefined behavioural repertoire, an alternative approach is to quantify the statistical dependence between animals by measuring whether knowledge of one animal’s location can reduce uncertainty about where another animal is located.

Information theory provides a framework for quantifying such statistical relationships without requiring specific interaction types to be defined a priori. Mutual information (MI) measures the amount of information that one variable provides about another and can therefore be used to quantify statistical dependence between the spatial trajectories of co-housed animals.^25^ In this context, higher MI indicates that an individual’s location can be more reliably predicted from knowledge of the locations of its cagemates, providing a measure of coordinated spatial behaviour independent of pre-defined behaviours. MI can further be used to characterize preferred configurations and higher-order interactions within groups of mice, capturing relationships that are not readily represented in conventional pairwise measures.^26^ Therefore, information theory offers a useful framework for quantifying relationships between animals from longitudinal home-cage positional data.

However, statistical dependence alone does not necessarily imply social interaction. In a bounded environment such as the traditional housing cages, animals are constrained by the same physical space, and their trajectories may become statistically dependent simply because they share environmental boundaries or spatial preferences. Thus, two animals moving independently within a small number of available locations can still exhibit measurable MI even in the absence of social influence. A meaningful measure of social dependence therefore requires comparison with a null model that preserves the relevant characteristics of individual movement within a cage while disrupting the dependence between cagemates. This baseline allows dependence arising from shared environmental constraints to be distinguished from dependence associated with social coordination.

To test whether statistical dependence between co-housed animals reflects meaningful social behaviour, we address three important questions in our validation of social behaviour within the home cage. First, can statistical dependence between co-housed animals be measured reliably? Second, what level of dependence is expected in the absence of social interaction? Third, does the resulting signal correspond to known strain differences and directly observed social behaviours? Comparing strains with established differences in sociability and independently observed social interactions provides additional validation of the relevance of this measure, while longitudinal monitoring allows its reproducibility and reliability to be assessed over time.

In this study, we developed a computational approach that leverages positional data from co-housed mice within their home cage to quantify social dependence using information theory. We focused on C57BL/6J and BTBR mice, two strains with well-established differences in sociability, as high- and low-sociability reference strains, respectively.^27^ We first evaluated whether MI distinguished observed home-cage behaviour from simulated null interactions and examined the sensitivity of this measure to temporal sampling parameters. We then assessed strain differences, reproducibility across repeated periods of home-cage monitoring, and correspondence between MI and independently observed social behaviours using video scoring. We further compared home-cage measures with sociability assessed using the canonical three-chamber task and examined relationships between brain and behavioural measures. Together, these analyses establish and validate an information-theoretic framework for quantifying social dependence from longitudinal home-cage monitoring and provides a basis for investigating the temporal dynamics and neuroanatomical correlates of social behaviour.

## RESULTS

Mouse behaviour was monitored continuously in standard cages using the Actual Analytics Home Cage Analyser (Actual Analytics, Edinburgh, UK), which tracks RFID-implanted mice across 18 antenna sensors arranged in a two-dimensional grid beneath the cage floor (**Figure 1**). This configuration provides the ability to longitudinally track each mouse’s position, allowing spatial occupancy and movement patterns to be quantified. These positional data provide the basis for deriving information-theoretic measures of social behaviour that captures statistical dependences between movements of mice and their cagemates, rather than relying on spatial proximity alone.

**Figure 1:**
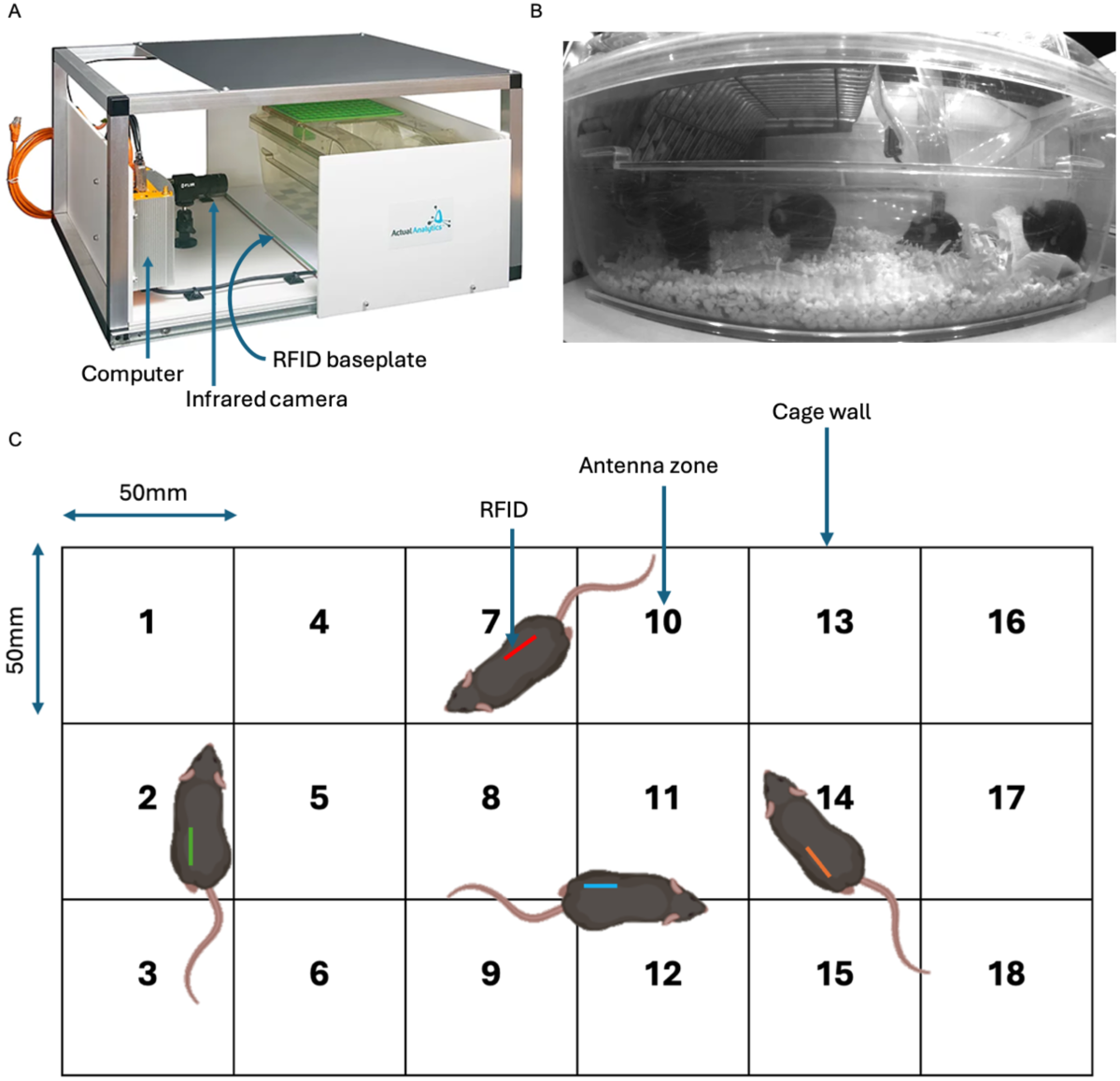
Home-Cage analysis mouse system. **A)** Single desktop unit consists of an infrared camera, computer, RFID baseplate array, acrylic lighting backdrop. The full mouse rack can fit multiple units which allows for simultaneous recording of up to 8 cages. **B)** Representative side-view snapshot of the home cage. Mice were group-housed in standard IVC cages with food, water via a sipper, bedding and enrichment. Videos were captured from the camera positioned on the long edge of the cage. **C)** Representative top-down view of the baseplate that tracks continuous activity in the home cage. The numbers correspond to the RFID antenna receivers on the baseplates used for calibration and detection of mouse position. The distance between each antenna is roughly 50mm.

To illustrate this approach, **Figure 2** shows how the spatial distribution of one mouse’s position can change conditional on the location of a cage mate. A shift in its positional distribution relative to the mouse’s typical distribution indicates that its behaviour is dependent on the cage mate’s location. Information-theoretic measures quantify these dependencies, providing a framework for evaluating social behaviour while accounting for individual spatial occupancy patterns. However, such dependencies do not necessarily reflect direct social interactions, as they may also arise from shared environmental preferences or other factors. We therefore implemented a null model to tease apart statistical dependence that reflects social behaviour compared to non-social influences.

**Figure 2:**
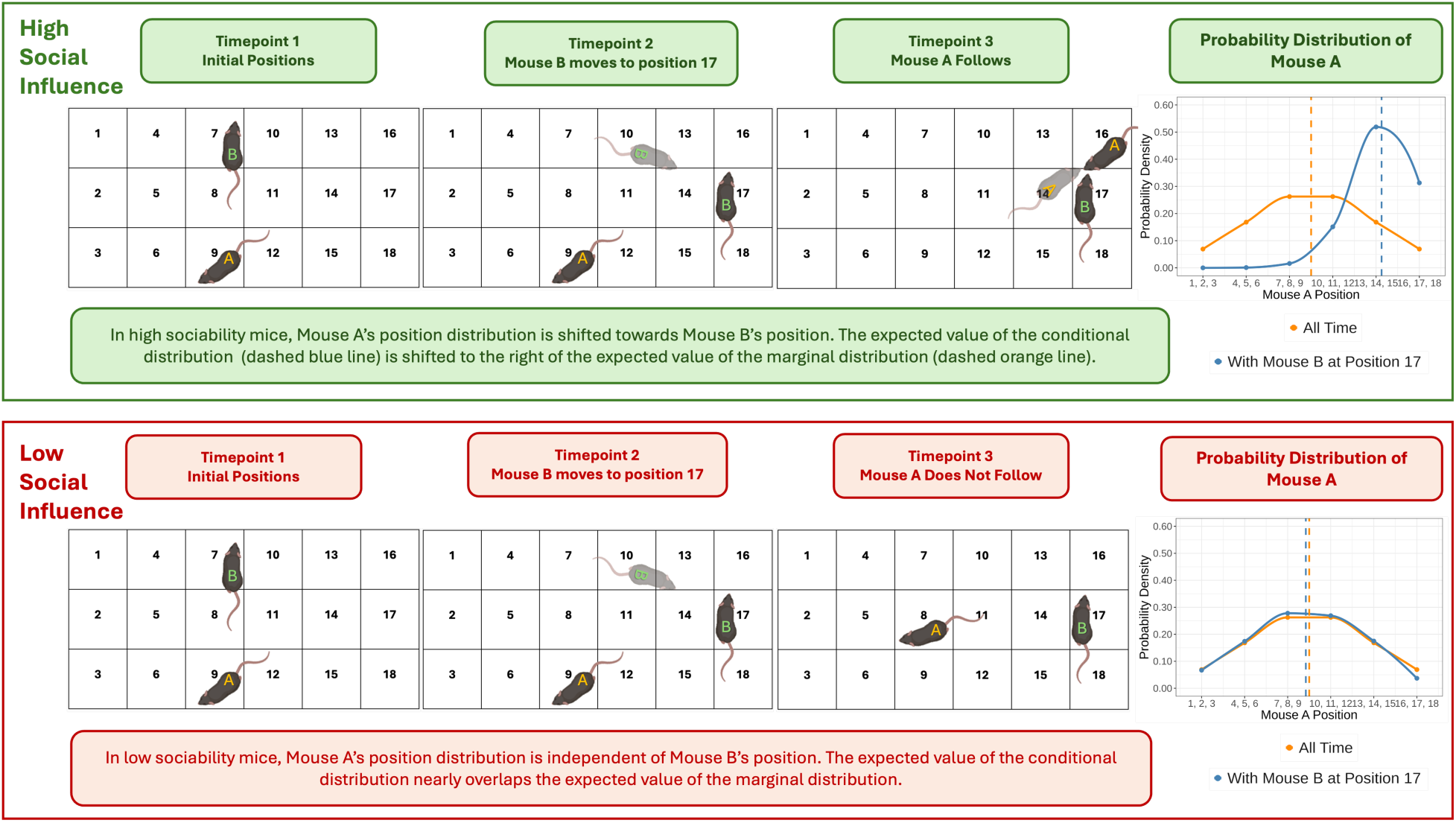
Probability distribution of a representative mouse is affected by a cage mate. The probability is measured at each antenna (points) and interpolated (lines). The orange points and orange solid line show the probabilities and distribution of Mouse A over the course of the night. The dotted orange line indicates the expected value of Mouse A’s position without any influence from its neighbours. The blue points and solid blue line show the conditional probabilities and distribution of Mouse A’s position when its cage mate Mouse B is located at antenna position 17. Higher values indicate higher probability of a mouse being in that position. Note the shift of the dotted blue line indicating the expected value of the conditional distribution is to the right of the dotted orange line, indicating a preferential shift in Mouse A’s position distribution towards position 17 where Mouse B is.

### Validation of mutual information as a measure of home-cage social behavior

We first validated MI as a measure of home-cage social behaviour using a constrained null model, established its reproducibility, and tested its correspondence with known strain differences and directly observed social interactions.

#### A constrained null model distinguishes social dependence from spatially induced mutual information

MI quantifies statistical dependence between the spatial positions of two mice, but non-zero MI or low levels of dependence can arise even in the absence of social behaviour because of the shared and spatially constrained environment. To distinguish genuine social dependence from dependence arising from shared spatial constraints, we compared the observed MI computed between mice housed together in the same cage with MI computed from the movement trajectories of mice randomly selected from different cages. This constrained null model preserved strain-specific movement characteristics while disrupting social dependence between cage mates. We evaluated real and null MI across four temporal bin and lag durations (30s, 60s, 300s, 900s) and identified the parameter combination that maximized separation between the two distributions.

Short aggregation windows (30s and 60s) frequently produced a high proportion of zero and near-zero MI values because mice were often stationary, resulting in limited variation in position occupancy and sparse sampling across antenna locations. Consequently, MI estimates at shorter timescales were more variable and susceptible to noise, likely reflecting inactivity rather than true social interactions. Across all temporal windows, real and null MI distributions showed substantial overlap at the individual-mouse level (**Figure 3A**). In contrast, separation between real and null data emerged following cage-level averaging, with observed cages exhibiting higher MI than their corresponding constrained null distributions (**Figure 3B**).

**Figure 3:**
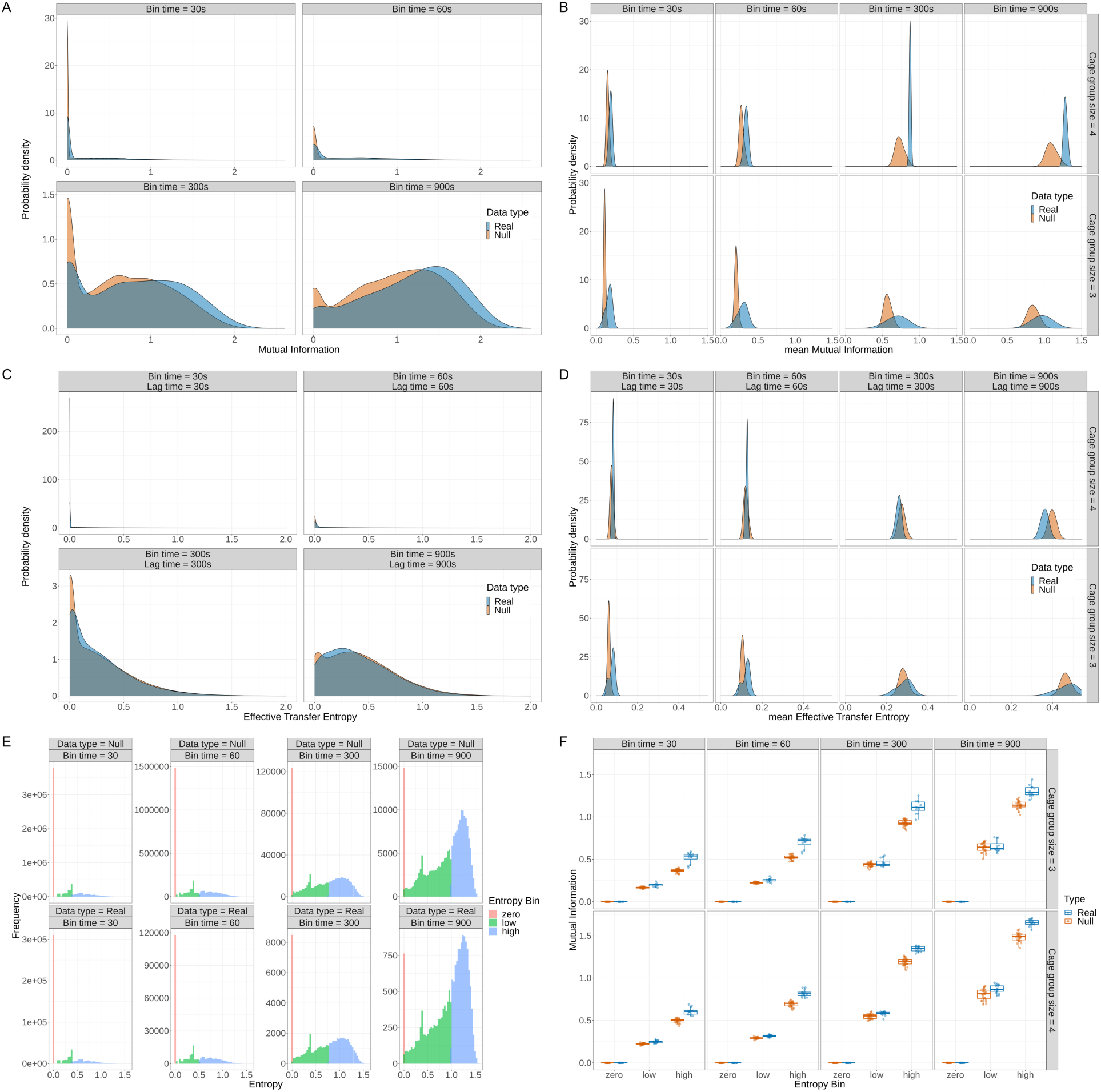
Validation of MI and TE against the constrained null model in C57BL/6J mice. **A)** Probability density plots of real distribution compared to null distribution averaged per mouse. Kernel density estimates are shown for real data (blue) and null data (orange), computed using bin durations of 30s, 60s, 300s, and 900s (columns). Increased zero and near-zero values observed at smaller time bins of 30s and 60s. **B)** Distribution of mean mutual information (MI) values for real versus null cages across multiple temporal bin sizes and cage group sizes (top row: 4 mice; bottom row: 3 mice). Across all bin times and group sizes, real cages consistently exhibit higher MI distributions relative to null. **C)** Distribution of mean effective transfer entropy (TE) computed across four different time bins averaged across mice and **D)** across cages. Only effective TE is shown since the affective TE produced identical values when averaged at the cage level. Directional differences between the two types of TE were only observed at the individual mouse level and are therefore not represented here. **E)** Histograms showing the distribution of entropy values calculated using temporal bin widths of 30, 60, 300, and 900 seconds for both data types within the C57BL/6J strain. Entropy values were classified as zero (pink), low (green), or high (blue), with low and high entropy defined relative to the median of non-zero entropy values for each bin width. Both data types exhibited a large proportion of zero-entropy windows at shorter time bins, while longer time bins resulted in a greater proportion of high-entropy states. Null distribution exhibited the same pattern, but at higher frequency given data was computed for each mouse and thus no random pairing. **F)** Boxplots represent mean MI per mouse (points) across entropy bins for real and null datasets in both cage sizes of C57BL/6J mice. Real datasets exhibited substantially higher MI than null data across entropy bins, particularly at high entropy bins.

For each combination of temporal window and group size, we tested whether mean cage-level MI in real cages exceeded that expected by chance, using permutation-based p-values computed from the null distribution (Benjamini-Hochberg (BH)-adjusted). Z-scores are reported as effect sizes only. In C57BL/6J, cage-level MI in real cages was higher than in the strain-matched null at all four temporal windows and for both group sizes (permutation-based p < 0.0001 for all eight comparisons, BH-adjusted; Z = 1.95-5.10, effect size; **Supplementary Table 1**). Cages containing three mice showed a broader range of MI values than four-mouse cages, whereas MI values in four-mouse cages were more tightly distributed (**Figure 3B**). The current time bins were selected to maintain the temporal resolution required for fine-scale behavioral analysis. Of note is since these temporal windows may under sample the large joint state space, the null distribution MI increases with bin size is likely due to finite-sample estimator bias. While infinitely large bins may allow the null MI to approach its true baseline, doing so would sacrifice the smaller-scale temporal dynamics needed for capturing real-time social interactions. As such, we selected a sampling time window of 300 s (5 min), which provided a balance between preserving temporal resolution and producing robust deviations from null expectations across both strains for downstream analyses of home-cage social metrics.

Since MI is bounded by the entropy of the underlying position distributions, we further examined the relationship between MI and entropy within a cage. Entropy was divided into zero, low, and high states, representing increasing levels of spatial exploration and behavioural diversity (**Figure 3E**). Zero entropy occurred more frequently at shorter aggregation windows, consistent with periods of limited movement. Under these conditions, MI was necessarily close to zero because there was little available positional information from which statistical dependence could be detected. Across both cage group sizes, as entropy increased, observed MI increasingly diverged from the constrained-null distribution, demonstrating that the distinction between social dependence and chance spatial co-occurrence was most apparent when sufficient movement was present (**Figure 3F**).

We performed similar analyses for transfer entropy (TE). Unlike MI, TE did not consistently distinguish observed cages from the corresponding constrained null distributions across temporal windows (**Figure 3C, D**). Real cage TE exceeded the null only in three-mouse cages at the two shortest time windows (30 s: Z = 2.79, p < 0.0001 BH-adjusted; 60 s: Z = 1.90, p = 0.0003 BH-adjusted), whereas four-mouse cages did not differ from the null at these windows (p ≥ 0.087 BH-adjusted). At longer time windows, three-mouse cages did not differ from the null (p = 0.67 BH-adjusted), and four-mouse cages showed TE significantly *below* the null (300 s: Z = −1.08, p = 0.044; 900 s: Z = −2.37, p < 0.0001, BH-adjusted). Extending the analysis across a broader range of temporal bins and lags similarly failed to identify a consistent, biologically interpretable lag structure (**Supplementary Figure 1**). Together, these findings indicated that MI provided a more robust measure of cage-level social dependence than TE and was therefore used for subsequent analyses.

#### Mutual information recapitulates known strain differences in social behaviour

Having established that observed MI exceeded expectations from a constrained null model in C57BL/6J mice, we next examined whether this pattern generalized across strains and whether MI could recapitulate a well-characterized difference in social behaviour between BTBR and C57BL/6J mice.^27^ Across both strains, observed MI exceeded null MI in both strains (**Figure 4A**), further supporting the conclusion that the measured dependence between cagemates’ spatial behaviour is not explained solely by strain-specific movement patterns and spatial constraints.

**Figure 4:**
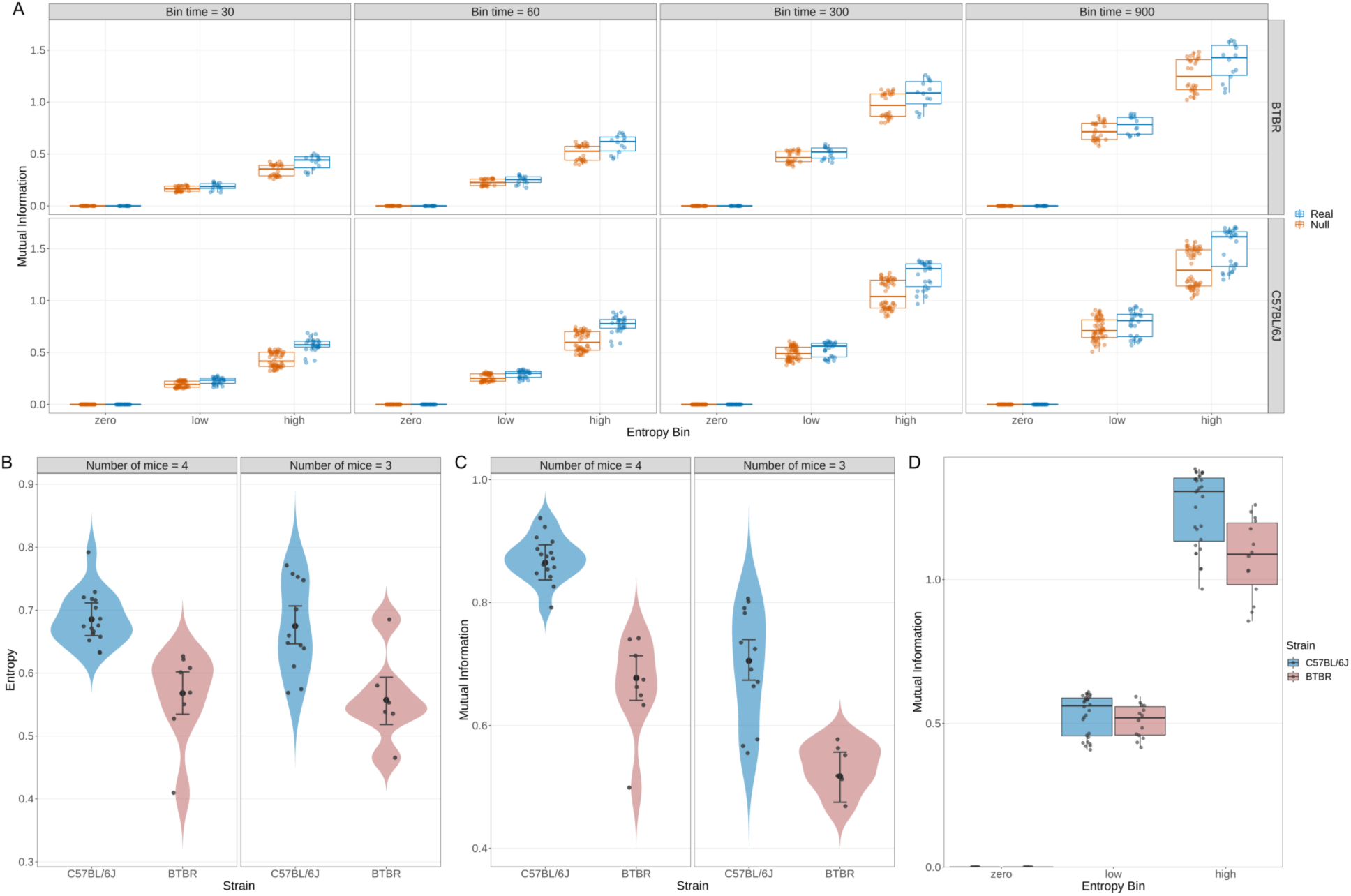
Home-cage entropy and MI recapitulate strain differences in social behaviour between C57BL/6J and BTBR mice. BTBR mice exhibited lower entropy and MI than C57BL/6J mice, with the strain difference in MI most pronounced at high entropy. **A)** Boxplot comparison of MI between real cage mates and strain-matched null pairs across both strains, demonstrating higher MI in real than null data. **B)** Violin plots showing mean entropy and 95% C.I. across all collected data filtered for the night phase. Data points represent average entropy per mouse. Two linear mixed-effects models were fit to the data. The first had main effects of strain, number of mice, and a random intercept for each mouse, and the second model had an interaction effect between strain and number of mice. The first model was used to determine the mean and 95% confidence intervals. After fitting both models, a likelihood ratio test was conducted and showed no significant interaction effect of strain and number of mice. **C)** Violin plot shows mean MI across all data collected, computed the same as entropy. Average mouse MI is represented as points and C.I. represent 95% of the bootstrap. **D)** Boxplot showing MI for BTBR and C57BL/6J mice across zero, low, and high entropy states. Points represent average MI per mouse. Low and high entropy were defined relative to the median of non-zero entropy values. Strain-by-entropy interactions were assessed using a linear mixed-effects model, followed by post hoc comparisons of estimated marginal means between strains within each entropy state. *P*-values for post hoc comparisons were adjusted across the three entropy states using the Holm method.

Since MI is influenced by the amount of spatial information available, we first examined entropy as a measure of spatial exploration. Low entropy indicates that the mouse position is stationary and visited few antennas, while high entropy would indicate the mouse position is relatively spread out and visited many antennas during the recording period, suggesting high exploratory behaviour and social curiosity. Entropy differed significantly between strains (p < 0.001), with BTBR mice exhibiting lower entropy than C57BL/6J mice across both three- and four-mouse cages (**Figure 4B**; Δ= 0.119, 95% CI [0.080, 0.158], p <0.0001). Consistent with this result, time-series analysis across the two-week recording period showed lower average entropy in BTBR mice (Supplementary Figure 2A). These movement patterns are consistent with BTBR’s characteristic repetitive behaviors,^11,27^ whereas C57BL/6J mice display more diverse exploratory behavior.

MI also differed significantly between strains (**Figure 4C**; p < 10^-11^). Post hoc contrasts via estimated marginal means revealed BTBR mice exhibited lower MI than C57BL/6J mice in both cage group sizes (Δ= 0.189, 95% CI [0.147, 0.231], p < 0.0001), with the same pattern observed across the time series data (Supplementary Figure 2B). Thus, MI recapitulated the established strain difference in social behaviour.

To determine whether strain differences in MI were attributable solely to differences in overall spatial exploration, we examined MI across entropy states. MI increased with entropy in both strains, but the magnitude of this increase differed between strains, as indicated by significant strain-by-entropy interactions at low entropy (β = −0.018, *p* = 0.007) and high entropy (β = −0.168, *p* < 0.0001). Relative to the zero-entropy reference state, BTBR mice showed a smaller increase in MI than C57BL/6J mice at both low and high entropy, with a greater difference at high entropy. Post hoc comparisons of estimated marginal means revealed no significant strain differences in MI at zero entropy (Δ = 0.008, 95% CI [−0.020, 0.037], Holm-adjusted p = 0.483) or low entropy (Δ = 0.023, 95% CI [−0.004, 0.049], Holm-adjusted p = 0.076). At high entropy, BTBR mice exhibited significantly lower MI than C57BL/6J mice (Δ = 0.175, 95% CI [0.149, 0.202], Holm-adjusted p < 0.0001; **Figure 4D**).

#### Home-cage mutual information shows high test-retest reliability and correlates with video-based measures of social behaviour

We next assessed the reproducibility of MI across the 14 recorded nights. Single measure intraclass correlation coefficients (ICCs) indicated moderate but significant reliability in both strains, with ICCs of 0.43 (95% CI [0.26, 0.68]) for BTBR and 0.45 (95% CI [0.32, 0.62]) for C57BL/6J. To visualize consistency across subsets of nights, we compared each mouse’s mean MI on odd- and even-numbered recorded nights (**Figure 5A**). The greater apparent agreement when averaging across multiple nights in the figure reflects the expected improvement in reliability obtained by averaging repeated measurements. These results demonstrate that individual differences in home-cage MI were reproducible across repeated nights of recording. To directly assess whether MI corresponded to observable social behaviour, we manually scored 5-min video segments selected from a range of MI values in BTBR and C57BL/6J cages. Segments represented low, medium, and high MI states and were centred on the corresponding MI time point. Social interactions, including sniffing, allogrooming, chasing, following, and mounting, were manually counted and summed to calculate a total social behaviour frequency. To minimize observer bias, video filenames were randomized and replaced with anonymized identifiers prior to scoring, such that the scorer was blinded to the corresponding MI values.

**Figure 5:**
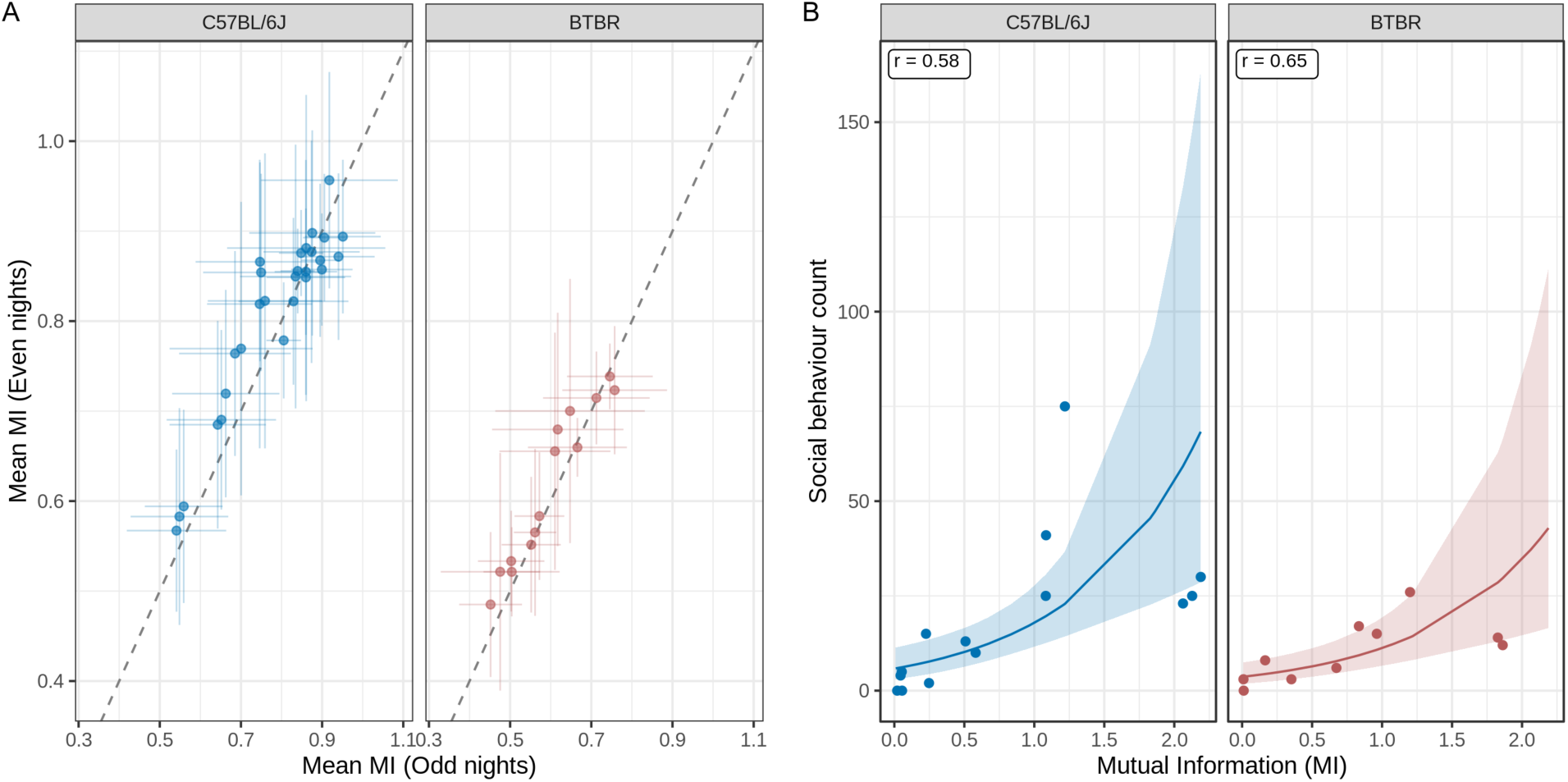
Validation of home-cage MI metric as a measure of social behaviour. **A)** Test-retest reliability of MI by strain. Each point represents a mouse, with mean MI on odd-numbered recorded nights plotted against mean MI on even-numbered recorded nights. Error bars represent standard deviation across the seven MI measurements. The dashed line indicates perfect agreement (y=x). The plot reflects agreement between seven-night averages rather than the reliability of a single night measurement. Formal test-retest reliability was quantified using one-way, single-measure ICC computed across all 14 nightly measurements per mouse. **B)** Correlation plot MI with observer video scoring of social behaviour. A total of 24 video clips (10 BTBR, 14 BL6/J three-mice cages) with a range of MI values were selected. Each point represents a 5-minute clip where social behaviour was scored. Both strains exhibit significantly strong positive correlation between MI scores and observer scores.

MI was positively associated with the number of socially interactive behaviours observed in recorded video clips (**Figure 5B**). Within each strain, MI was positively correlated with behaviour counts in both BTBR (r = 0.65, p = 0.040) and C57BL/6J (r = 0.58, p = 0.029) mice. To account for the count nature of the data and the repeated sampling of cages, we fit a negative binomial mixed model with cage as a random intercept. Each 1-unit increase in MI was associated with a 3.1-fold increase in expected behaviour count (rate ratio = 3.1, 95% CI 1.7-5.5, p < 0.001), after accounting for strain. Strain did not significantly affect behaviour count (BTBR vs C57BL/6J: rate ratio = 0.63, 95% CI 0.32-1.24, p = 0.18), and the MI by strain interaction was not significant (likelihood ratio test, χ² (1) = 0.08, p = 0.78), so a common MI slope was used for both strains. The estimated cage-level variance was near zero, indicating that clips from the same cage were not more similar than clips from different cages after accounting for MI and strain. Thus, clips with higher MI contained more directly observed social behaviour in both strains, providing further validation that MI captures meaningful social behaviour. Representative video segments corresponding to low and high MI values are provided in Supplementary Material.

### Application of MI to characterize home-cage social behaviour

Having established that MI exceeded movement-constrained null expectations and is a meaningful metric that is able to measure known differences in social behaviour, we next applied MI to characterize variation in home-cage social behaviour across genetic background, behavioural context, neuroanatomy, and circadian cycle. Additionally, we tested the extent with which MI and the three-chamber task capture similar or distinct aspects of social behaviour.

#### Home-cage social behaviour varies across genetic background and is distinct from three-chamber sociability

To determine whether the relationship between entropy state and MI observed in the BTBR and C57BL/6J comparison generalized across genetic backgrounds, we extended the analysis to seven inbred strains (C57BL/6N, C57BL/6J, BALB/cJ, DBA/2J, BTBR, FVB/NJ, A/J). A linear mixed-effects model including strain, entropy state, and their interaction revealed strain-dependent differences in MI across entropy levels (Supplementary Figure 3). MI differences were negligible at zero entropy and modest in the low entropy state, whereas pronounced strain-dependent divergence emerged at high entropy, with C57BL/6J mice exhibiting the largest increase in MI relative to other strains. These findings indicate that home-cage social coordination varies substantially across genetic backgrounds, particularly during periods of active exploration.

We next asked whether home-cage MI was related to sociability measured in the three-chamber task. The three-chamber sociability index (SI) reflects preference for proximity to a novel conspecific, whereas MI quantifies statistical dependence between familiar cage mates. Since both measures reflect unique but related aspects of social engagement and behavioural interaction, we hypothesized that they would capture overlapping features of social behaviour and therefore exhibit some degree of correlation. Interestingly, no significant correlation was observed between home-cage MI and the three-chamber SI, indicating that dependence between cage mates in the home cage was not associated with time spent investigating the novel mouse in the three-chamber task. (**Figure 6**; *r* = 0.07, p = 0.879).

**Figure 6:**
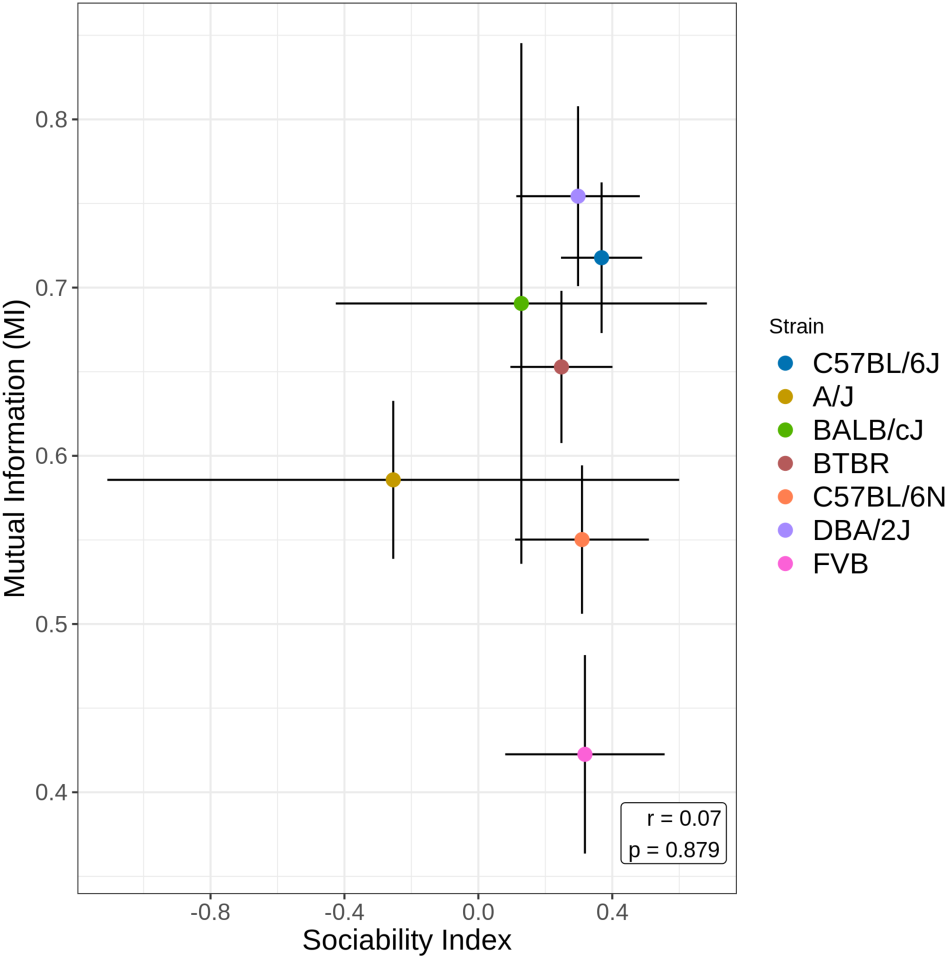
Strain differences in sociability across seven strains. Correlation between home-cage MI is plotted against three-chamber sociability index (SI). SI is defined as the difference of time spent with stranger and empty cup over sum of time with stranger and cup. Points represent strain averages. Error bars represent 95% confidence intervals in SI (horizontal) and MI (vertical). No significant correlation was found (r = 0.07, p = 0.879).

#### Home-cage social behaviour has distinct neuroanatomical associations

Given the absence of a significant association between HCM-derived MI and the canonical three-chamber social behaviour measure, it raises the question of whether HCM-derived MI nevertheless captures biologically meaningful variation. Thus, we next examined the neuroanatomical correlates of home-cage MI using *in vivo* magnetic resonance imaging (MRI). Structural MRI provided an independent means of examining whether strain differences in brain volume involved regions previously implicated in social behaviour, and whether neuroanatomical correlates differed across behavioural measures derived from home-cage monitoring, the three-chamber test, and the open field test (OFT).

Regional brain volumes were normalized to whole-brain volume, and Pearson correlations were calculated between regional volume and MI, home-cage entropy, and three-chamber SI. Multiple comparisons were controlled using a Benjamini-Hochberg false discovery rate (FDR) of 10%. Across all mice and without adjustment for strain, 26 brain regions showed significant associations with MI following FDR correction (**Figure 7A**). Significant positive correlations were observed in regions including the medial septal complex (*r* = 0.66, FDR-adjusted p = 7.97 x 10^-6^), lateral septal complex (*r* = 0.53, FDR-adjusted p = 1.47 x 10^-3^), and hypothalamus (*r* = 0.52, FDR-adjusted p = 1.85 x 10^-3^). Negative correlations were observed in several cerebellar regions, including the simple lobule (*r* = −0.53, FDR-adjusted p = 1.46 x 10^-3^) and declive (*r* = −0.51, FDR-adjusted p = 1.95 x 10^-3^), as well as the ventrolateral part of the orbital area (*r* = −0.56, FDR-adjusted p = 8.35 x 10^-4^).

**Figure 7:**
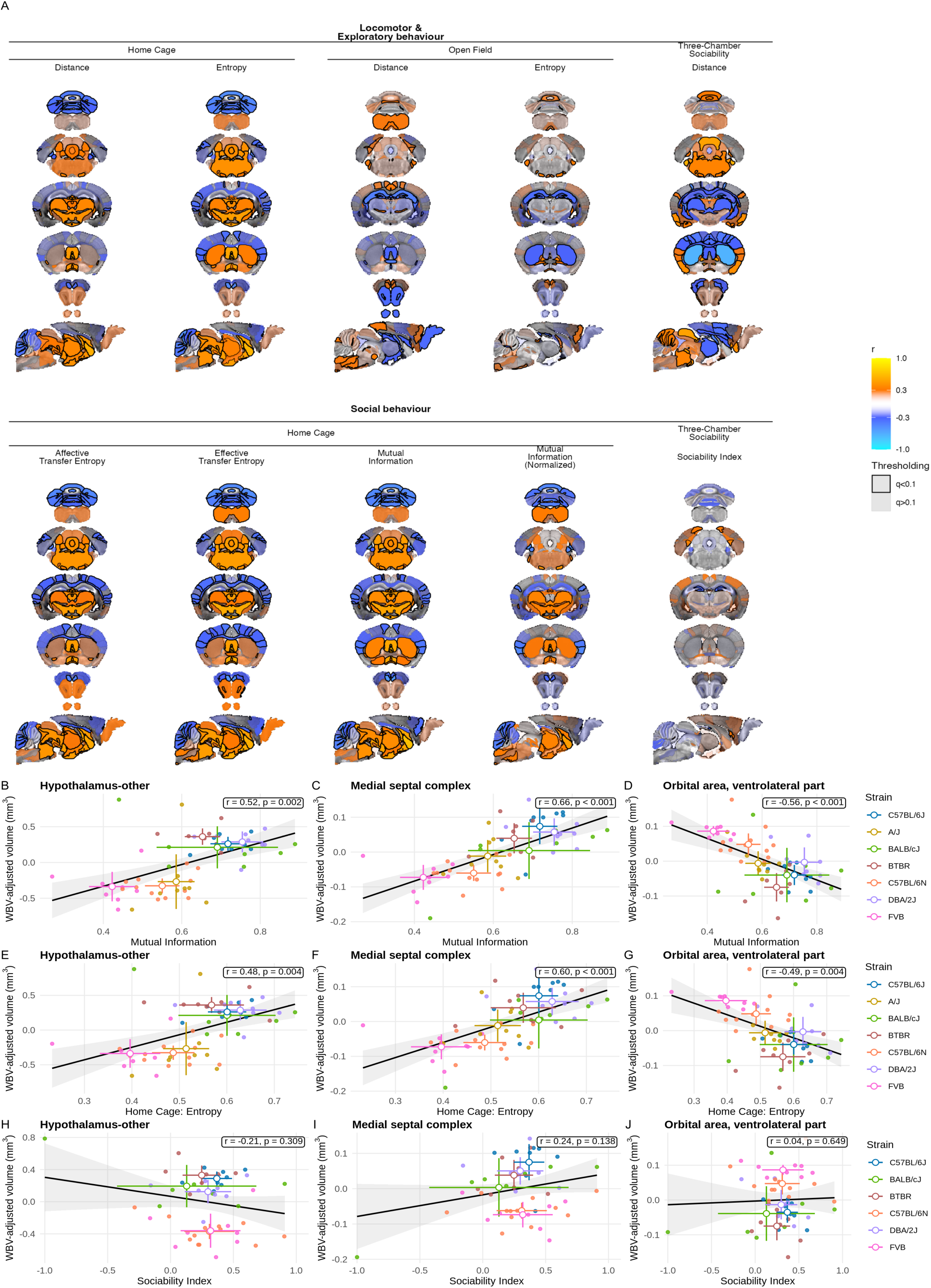
Correlations between brain structures and behaviour scores across all mice. A) Volumes per mouse were computed for every brain region and correlated against behaviour scores. Warmer orange/yellow colours indicate positive correlation, meaning as the specific behaviour score increases, so does brain volume, whereas cooler blue colours indicate negative correlation, meaning behaviour score increases are associated with brain volume decreases. Brain volumes were adjusted for overall brain volume. Structures that survive multiple comparisons (BH FDR 10%) are outlined in black. **B)** to **D)** show significant correlations between MI and normalized volumes of hypothalamus, medial septum, and ventral orbital cortex. **E)** to **G)** show significant correlations between entropy and the same three structures. **H)** to **J)** show insignificant correlations between SI in the three-chamber task across the three structures. Solid colored points represent a mouse within each strain and white points (with corresponding colour borders) represent strain averages. Cross error bars represent 95% C.I. for each strain and the gray band shows 95% confidence interval around the fitted regression line. Mice in the A/J strain were excluded from SI correlation given they remained largely stationary during the task.

Given MI is constrained by the entropy of the underlying position distribution, we next compared its neuroanatomical associations with those of home-cage entropy. Substantial overlap was observed between regions associated with MI and entropy, consistent with the strong correlation between these measures. However, seven structures were significantly correlated with MI but not entropy: Culmen, insular region, nucleus accumbens, optic tract, medial part of the orbital area, trunk region of the primary somatosensory cortex, and the superior olivary complex (Supplementary Table 2). Normalizing MI by entropy further identified significant correlations in additional structures including Crus 1, fastigial nucleus, interposed nucleus, cerebellar lobules, and the medulla. Together, these findings suggest that, despite shared neuroanatomical associations with spatial entropy, MI additionally captured distinct patterns of brain-behaviour association that were not explained from entropy alone, suggesting a contribution of social behavioural information beyond general exploratory behaviour.

We next compared these associations with the three-chamber sociability. Four structures were moderately correlated with SI in the three-chamber task after multiple comparisons (Supplementary Table 2), including the subiculum (*r* = 0.35, FDR-adjusted p = 0.04), dentate gyrus (*r* = −0.35, FDR-adjusted p = 0.05), the posteromedial visual area (*r* = 0.35, FDR-adjusted p = 0.05), and the mediomedial area of the secondary visual cortex (*r* = 0.31, FDR-adjusted p = 0.09). Three regions of interest were chosen a priori based on their previously documented roles in regulating and modulating social behaviour in rodents.^28–30^ Comparing these regions of interest across all three metrics, we found similar trends in correlations with MI and entropy where the hypothalamus and medial septum both exhibited significant positive correlations (**Figure 7B, C, E, F**), while the ventral orbital cortex (**Figure 7D, G**) was negatively correlated with both home-cage metrics. In comparison with the three-chamber task, none of the three structures of interest were significantly correlated with SI (**Figure 7H-J**). A/J mice were excluded from the SI correlation analysis because they remained largely stationary throughout the task, which could bias comparisons of sociability given their limited or lack of engagement with the novel stimulus.

Finally, we examined whether the neuroanatomical associations of home-cage entropy were specific to the home-cage environment by comparing with entropy measured during the OFT, where mice are tested individually in a novel environment. In contrast to home-cage entropy, OFT entropy showed a distinct pattern of significant neuroanatomical correlates in 23 structures (**Figure 7A**). Positive correlations were observed in cerebellar structures like the flocculus (*r* = 0.32, FDR-adjusted p = 0.01) and simple lobule (*r* = 0.26, FDR-adjusted p = 0.07). In contrast, negative correlations were observed in Ammon’s horn (*r* = −0.35, FDR-adjusted p =0.01), dentate gyrus (*r* = −0.35, FDR-adjusted p = 0.01), and Caudoputamen (*r* = −0.29, p = 0.03). Together, these results revealed context-dependent neuroanatomical correlates of entropy, suggesting that although entropy reflects spatial exploration in both settings, its neuroanatomical associations in the open field may additionally reflect anxiety-related processes that are less prominent in a familiar home-cage environment.

#### Social coordination exhibits distinct temporal dynamics across the circadian cycle

We next examined whether strain differences in MI were explained solely by differences in overall locomotor activity or whether social coordination exhibited distinct temporal dynamics across the circadian cycle. Across the seven-strain cohort, substantial strain differences were observed in total distance travelled and antenna transitions during 5-min intervals. Relative to C57BL/6J mice, A/J, BALB/cJ, C57BL/6N, and FVB mice travelled significantly less distance, whereas BTBR and DBA/2J did not differ significantly from C57BL/6J (**Figure 8A**). Similar differences were observed for antenna transitions (**Figure 8B**), with the largest difference between C57BL/6J and FVB mice for both total distance travelled (Δ = 1218.0 mm, *p* < 0.0001) and antenna transitions (Δ = 295.7, *p* < 0.0001).

**Figure 8:**
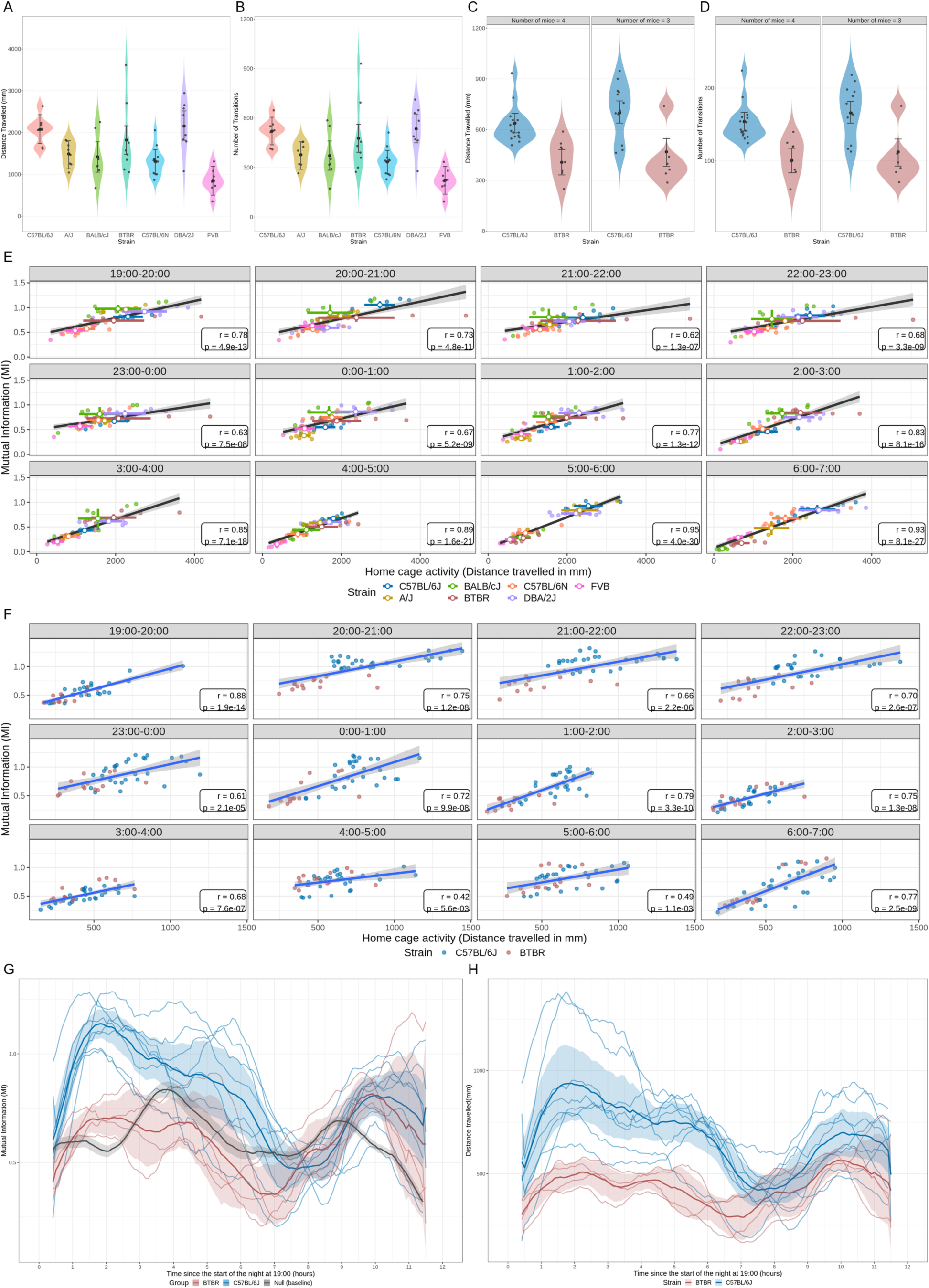
Strain differences in home-cage locomotor activity and MI. **A)** Strain differences in total distance travelled and **B)** total number of antenna transitions measured within each 5-min interval across the seven-strain cohort. A/J, BALB/cJ, C57BL/6N, and FVB mice exhibited significantly lower distance travelled and fewer antenna transitions relative to C57BL/6J, whereas BTBR and DBA/2J did not differ significantly from C57BL/6J. The largest strain differences were observed between C57BL/6J and FVB for both total distance travelled and antenna transitions. BTBR mice have significantly lower activity than C57BL/6J mice in **C)** Mean distance travelled and **D)** Mean number of transitions across all collected data filtered for the active period (night phase). **E)** Hourly correlations between distance travelled and MI across the mixed-strain cohort. Each solid point represents the mean distance travelled at each hour of the night which is plotted against mean MI for each mouse. White points represent strain averages with the respective strain colour around the border. Linear regression lines indicate overall trends within each hour. Pearson’s correlation (r) in each panel correspond to the association between activity and MI for that time window. Correlations remained significant throughout the night, with associations strengthening toward the end of the dark phase. **F)** Hourly correlations between distance travelled and MI in C57BL/6J and BTBR mice, with significant associations observed at the beginning, middle, and end of the night. **G)** Sliding window estimates of MI between mice are plotted as a function of time since the start of the dark phase (19:00) over a 12-hour period. For each 5-minute sampling time, MI was averaged over an hourly window centered on that timepoint, consisting of six preceding and six subsequent 5-minute bins each hour. Lines represent mean MI for each group (C57BL/6J, blue; BTBR, red; Null, grey), with shaded regions indicating 95% confidence interval. Thin lines show individual cage-level trajectories. **H)** Sliding window estimates of distance travelled averaged across one night cycle computed the same as MI.

In a separate BTBR and C57BL/6J cohort, we examined whether differences in MI were accompanied by differences in locomotor activity. During the active dark phase, BTBR mice exhibited lower distance travelled (Δ = 235 ± 43.8, *p* < 0.0001) and fewer antenna transitions (Δ = 54.3 ± 9.93, *p* < 0.0001) than C57BL/6J mice (**Figure 8C, D**). Despite these differences in overall activity, the relationship between activity and MI varied across the night.

Across the mixed-strain cohort, activity and MI remained significantly correlated throughout the night, but the strength of this association varied across the night (**Figure 8E**). Correlations were weaker at the beginning of the night (21:00–22:00: *r* = 0.62, p = 1.3 × 10^-7^) and strongest toward the end of the dark phase (5:00–6:00: *r* = 0.95, p = 4.0 × 10^-30^; 6:00–7:00: r = 0.93, p = 8.1 × 10^-27^), indicating that the coupling between locomotor activity and MI strengthens as the night progresses across strains. In a separate cohort of C57BL/6J and BTBR mice, activity and MI were also positively correlated throughout the night, but the strength of this relationship varied temporally, with the weakest association occurring at approximately 04:00 and 06:00, when the two strains showed greater convergence in MI and activity (**Figure 8F**). These findings demonstrate that despite reduced overall activity, BTBR mice exhibited a similar positive relationship between activity and MI throughout the night cycle, although the strength of the relationship fluctuated alongside the degree of strain separation at each hour.

To characterize the temporal organization of these measures more directly, we aligned two weeks of nighttime recordings to a common night start time (19:00) and calculated hourly sliding-window averages for each 5-min interval. In the cohort with only BTBR and C57BL/6J mice, BTBR mice exhibited lower MI and locomotor activity than C57BL/6J mice across most of the night. The strains converged during the late-night period, approximately nine hours after night onset, when BTBR MI increased toward that of C57BL/6J mice. Both MI and distance travelled exhibited a biphasic pattern across the dark period, with an initial increase following lights-off and a peak approximately two hours into the night, followed by a decline during the middle of the active period and a second, smaller increase approximately two hours before lights-on (**Figure 8G, H**). In contrast, this bimodal pattern was not observed during the inactive light phase (Supplementary Figure 4A, B). The mixed-strain cohort showed a similar pattern, although peak timing varied across strains (Supplementary Figure 4C, D).

Together, these findings indicate that home-cage social coordination and locomotor activity are temporally coupled but dynamically vary across the circadian cycle, and that differences in overall activity do not fully account for the temporal organization of MI.

## DISCUSSION

Social behaviour in rodents is dynamic, context-dependent, and expressed across timescales that are often difficult to capture using conventional behavioural assays. Automated home-cage monitoring (HCM) complements these approaches by enabling continuous measurement of spontaneous behaviour in familiar, group-housed environments, while reducing experimenter intervention and exposure to novel testing environments.^31,32^ HCM has increasingly been used to characterize strain- and disease-related phenotypes, including social behaviour, in freely interacting animals.^31,32^ Here, we developed an information-theoretic framework that uses spatial occupancy to quantify the statistical dependence between cagemates as a measure of social behaviour. Across longitudinal home-cage recordings, MI distinguished observed cages from constrained null distributions, recapitulated known social differences in BTBR compared to C57BL/6J mice, showed moderate but significant reproducibility across repeated nights, and corresponded to direct observations of social interactions. Using this framework, we demonstrated that home-cage social behavior varied across genetic background, has neuroanatomical associations that are distinct from general spatial exploration and three-chamber sociability, and exhibited temporal organization across the circadian cycle. Rather than replacing currently established behavioural assays, our approach builds on the growing use of automated home-cage monitoring to capture complex behavioral phenotypes over extended periods.^31,32^

A central challenge in using MI to quantify social behaviour is distinguishing statistical dependence arising from non-social factors from that associated with social interaction. Although the physical constraints of the home cage limit the variability in positional configurations, these spatial constraints alone are insufficient to generate MI. Systemic temporal variation in activity and spatial occupancy may produce apparent MI even in the absence of direct social interaction. For example, mice in different cages may exhibit similar movement patterns at certain times of the night, resulting in statistical dependence. Our constrained null model was designed to provide an estimate of the MI that can arise under these conditions by comparing mice from different cages while preserving strain-specific movement, individual positional distributions, and temporal sampling characteristics from the observed data. Observed cage-level MI consistently exceeded these constrained-null expectations, suggesting that co-housed mice exhibit greater statistical dependence beyond what would be expected from individual movement patterns and the associated temporal structure alone.

The choice of temporal resolution in our study was also important as it required balancing the ability to capture fine-scale behavioural dynamics against the stability of information-theoretic estimates. We evaluated the influence of temporal parameters on information-theoretic metrics by systematically varying bin duration and time-lag parameters, comparing the resulting distributions between real and null data at both the individual mouse and cage levels. At very short timescales (30 and 60 s), mice were frequently stationary, resulting in limited variation in position occupancy and a high proportion of zero or near-zero MI estimates. Consequently, these estimates may reflect insufficient positional variation rather than an absence of social dependence. At intermediate timescales, including the 300s windows used in our subsequent analyses, MI increased in both real and null data. One possible explanation for this trend is finite-sample bias in discrete MI estimation. The joint position state space grows exponentially with the number of mice, encompassing up to 18^n^ possible joint states for a group of n mice. Since only a subset of these states may be observed within a given sampling window, estimates of the joint and marginal probability distributions may be affected by sparse sampling. Under the null hypothesis, random fluctuations in these estimated probabilities can produce positive MI estimates even when the underlying variables are independent, potentially contributing to the observed increase in null MI with longer bin durations. However, the extent to which estimator bias explains this trend depends on the effective sample size and the implementation of temporal aggregation. Although longer windows capture behaviour over extended periods, they do not necessarily reduce finite-sample bias, particularly when the joint state space is large relative to the number of observations. Furthermore, longer windows reduce temporal resolution and may obscure the fine-scale dynamics of social interactions that home-cage monitoring is intended to capture.

To assess the potential influence of finite-sample bias on MI estimates, we also compared our results with that obtained using shrinkage entropy estimators. The shrinkage estimator provided an alternative approach to estimating the underlying probability distributions, allowing us to evaluate whether our findings were sensitive to the choice of estimator. MI estimates obtained using the two methods were similar when calculated from the same observations. Importantly, the central finding that MI was greater among real cagemates than among strain-matched null distribution was maintained across all temporal windows and cage sizes using both estimators. Together, these findings support the robustness of our primary observed MI estimates to the choice of entropy estimator, while highlighting the importance of considering estimator behaviour when analysing sparsely sampled joint state spaces.

Although both MI and TE can be used to characterize statistical relationships within the home cage, our findings showed MI, compared to TE, provided a more robust measure of social dependence under the conditions examined here. MI consistently distinguished observed cages from their constrained-null distributions, whereas TE showed less consistent separation across temporal windows and cage sizes. This difference is consistent with an important distinction between the two measures: whereas MI quantifies statistical dependence between variables without requiring an explicit temporal direction, TE is defined by the conditional dependence of a future state on the past of another variable and therefore requires assumptions about the relevant temporal structure of the interaction.^33^ Although TE can provide information about directional relationships between animals, its interpretation depends on selecting parameters that appropriately capture the underlying behavioural process.^34^ In behavioural studies, this presents a particular challenge because the relevant timescale of an interaction may not be known a priori. Very short time lags may fail to capture interactions occurring over longer behavioural timescales, whereas long lags can produce apparent information transfer arising from coincidental temporal structure rather than genuine social interaction. Our analysis of TE across a broad range of aggregation windows and temporal lags did not reveal a consistent or biologically interpretable signal. Instead, elevated TE values occurred across a broad range of time lags, including delays that were unlikely to represent direct mouse-to-mouse interactions. Thus, in the present application, the absence of a clear temporal structure favoured the use of MI as a more robust measure of social dependence, while TE may be more informative when the temporal structure of the underlying interaction is sufficiently well characterized to guide lag selection and when recordings provide sufficient observations to estimate the relevant conditional probability distributions reliably.^34^

Our approach builds on previous applications of information theory to quantify social interactions in the home cage. In particular, Shemesh et al. demonstrated group behavior in mice cannot be explained by individual traits or pairwise interactions alone, requiring higher order (three-way) social interactions to accurately model group dynamics.^26^ We extend this framework by explicitly evaluating the extent to which observed MI exceeds statistical dependence expected from shared spatial constraints and strain-specific movement patterns, and by validating MI against strains with known social differences and directly observed social interactions. Using the selected temporal parameters, we demonstrated BTBR mice exhibited lower MI with their cagemates than C57BL/6J mice, consistent with reduced coordination of spatial behaviour and the established social phenotype of this strain.^35^ BTBR mice also exhibited lower entropy, suggesting reduced variability in their spatial behaviour. This provides a strong basis for interpreting information-theoretic dependence as a behavioural phenotype rather than simply as statistical association between animals sharing an environment.

The lack of correspondence between home-cage MI and three-chamber SI suggests that these assays capture different aspects of social behaviour. The three-chamber task primarily assesses social approach and is best interpreted as a qualitative measure of preference for a social stimulus rather than a quantitative measure of social interaction. Although social mice generally show a preference for a novel conspecific, the absolute amount of time spent with the stimulus can vary across studies and is a relatively coarse measure of social behaviour that may not capture the full repertoire of social interactions.^36^ The three-chamber assay is therefore well suited to assessing whether an animal demonstrates a preference for a social stimulus, whereas longitudinal HCM captures how animals interact within an established social group. This distinction is consistent with findings from Fairless et al., who reported that paired cage mates primarily engaged in passive social behaviors such as huddling and physical contact, whereas the introduction of a novel stimulus during the three-chamber assay elicited more active social exploration.^37^ The lack of significant correlation between assays should therefore not necessarily be interpreted as a limitation of either measure. Instead, it highlights the complex nature of social behaviour. A mouse may show a normal preference for a novel conspecific while exhibiting altered coordination with familiar cagemates, or vice versa. This distinction may be particularly relevant when studying neurodevelopmental and neuropsychiatric phenotypes, where social behaviour can be altered in specific contexts rather than uniformly across all forms of social interaction.

The longitudinal nature of HCM may additionally improve the stability of measures that are sensitive to behavioural context. In our study, MI showed moderate reproducibility across repeated nights and correlated with manually scored social interactions, supporting its use as a stable measure of spontaneous social behaviour. By comparison, Fairless et al. reported a weak test-retest correlation of r = 0.31 (p < 10^-4^) for repeated three-chamber testing in male and female C57BL/6J and BALB/cJ mice.^36^ These comparisons should be carefully interpreted since the assays measure different behaviours and were conducted under different experimental conditions. Nevertheless, they illustrate the advantage of repeated home-cage sampling for capturing stable individual differences and collective dynamics in social behaviour.

The neuroanatomical correlates provide further insight into the behavioural phenotypes captured within the home-cage. Raw MI was associated with several regions implicated in social behaviour, including the septal complex and hypothalamus, although many of these associations overlapped with those of home-cage entropy, consistent with the mathematical dependence of MI on positional entropy. Importantly, entropy-normalized MI revealed additional associations beyond those identified with raw MI, indicating that the relationship between cagemates’ behaviour captures information beyond spatial exploration. Together with the distinction between real and null MI and the recapitulation of strain differences in sociability, these findings support MI as a biologically meaningful measure of social behaviour that captures patterns of behavioural dependence not explained by exploratory behaviour alone.

Although entropy provides a common measure of spatial variability, its neuroanatomical correlates differed when measured in the home-cage compared to the open-field assay. Home-cage entropy was positively associated with the hypothalamus and septal complex, regions implicated in the regulation of social behaviour and social information processing.^30,38,39^ In contrast, open-field entropy was more strongly associated with hippocampal structures involved in spatial processing and the organization of exploratory behaviour.^40^ Associations with cerebellar structures in the OFT may reflect their established roles in sensorimotor and locomotor control of navigating a novel environment, whereas negative association with the hippocampus may reflect the contribution of emotional or anxiety-related responses to novelty.^41,42^ The different neuroanatomical correlates of entropy in the home cage and OFT demonstrate the importance of behavioural context in interpreting brain behaviour relationships. Although entropy quantifies spatial variability in both settings, home-cage entropy is measured in the presence of cagemates within a familiar environment and may therefore incorporate social influences. In contrast, open-field entropy reflects individual exploration in a novel, experimenter-administered environment, where anxiety-related responses may influence behaviour.

Furthermore, the neuroanatomical patterns associated with three-chamber SI were comparatively distinct from MI and home-cage entropy. These differences parallel the lack of correlation between MI and three-chamber sociability and provide convergent evidence that the two assays capture distinct aspects of social behaviour. Altogether, the neuroanatomical comparisons across MI, SI, and entropy suggest that home-cage monitoring captures a behavioural phenotype that is not reducible to general exploration and is distinct from social preference measured by the conventional three-chamber assay.

Finally, longitudinal monitoring within the home-cage revealed temporal organization of social dependence that would be difficult to capture using short-term acute assays. MI and locomotor activity showed similar early- and late-night increases, consistent with previous observations of increased exploration and social behaviour during the active phase.^43^ However, the strength of this relationship varied across the night, indicating that increased movement does not necessarily correspond to greater social dependence. Despite lower overall activity and MI, BTBR mice retained broadly similar nocturnal patterns in positional trajectories to C57BL/6J mice, suggesting that temporal organization of behaviour is largely preserved while its magnitude is reduced. While we did not directly assess circadian entrainment in this study, this pattern is consistent with previous evidence that BTBR mice remain entrained to environmental light-dark cycles despite alterations in intrinsic circadian properties.^44^ These findings further demonstrate the advantage of HCM in measuring when certain behavioural phenotypes emerge and how they may change across time.

### Limitations and future directions

There are a few limitations to the study that should be considered. Firstly, we examined only male mice. This choice reflects both practical considerations; easier to compare results with prior work and existing literature, as well as the known propensity for males in group housing to establish dominance hierarchies. However, sex is an important determinant of social behaviour, and prior studies have established sex differences in social structures, affiliative behaviors, and display of aggression.^45,46^ Future studies should therefore consider evaluating both sexes and determine whether the information-theoretic measures identified here generalize across sex.

Secondly, the present study focused on simple, non-antagonistic social interactions in adult mice, and we have yet to apply our approach to younger mice or investigate behaviors such as aggression and mating. Age would be particularly interesting to look at since there is evidence that social behaviours and interactions can change across development.^47^ Extending this framework across sex, age, genetic, and treatment models will help determine how broadly these measures capture social phenotypes and enable a more comprehensive characterization of normal and atypical social behaviour.

Although this study focused on examining the influence of other mice’s locations, persistent environmental factors, such as the placement of food and water, may also contribute to alterations in behaviours within the home cage. Under standard housing conditions, mice always have unrestricted access to food and water. However, in situations where these resources are limited, social dynamics may shift, potentially increasing competition and promoting the establishment of stronger hierarchical structures. While environmental variables like food and water availability were not explicitly considered in the current study, the methods presented here could readily be adapted to incorporate environmental and experimental manipulations such as enrichment or resource restriction.

Another important consideration is the challenge of disentangling entropy and social behaviour in a conventional home-cage environment. Although we attempted to tease apart “social” entropy from “exploratory” entropy by comparing home cage to open field metrics, since group-housed mice are housed in a relatively confined environment, an individual mouse’s movement is inherently influenced and restricted by the locations and movements of its cagemates. Consequently, home-cage entropy likely reflects both social and non-social components of behavioural variability, even when MI and entropy can be mathematically separated. This spatial constraint may also contribute to the overlap we observed between their neuroanatomical correlates. Environments that provide greater opportunities for exploration without spatial constraints, such as instrumented or enriched cages, may help separate variability arising from social interactions from that associated with non-social exploration.

## CONCLUSION

We developed and validated a constrained information-theoretic framework for quantifying social dependence from longitudinal home-cage trajectories. Such measures may therefore provide a means of quantifying continuous and subtle forms of social interaction that may not be apparent from conventional behavioural parameters. The framework establishes an empirically defined baseline for dependence expected from the physical and behavioural constraints of the cage, demonstrates that observed MI is reproducible and behaviourally meaningful, and provides a platform for discovering genetic, neuroanatomical and temporal variation in social behaviour.

## METHODS

### Mice

Two independent cohorts were used to address different study objectives while minimizing potential confounding effects of experimental procedures on home-cage behaviour. All mice were housed at The Center for Phenogenomics animal facility on a 12/12 h light/dark cycle with food and water *ad libitum*. Cage changes and food/water top ups were done once a week. Cohort one included seven inbred mouse strains and was used to investigate associations between home-cage social metrics, whole-brain structural magnetic resonance imaging, and conventional behavioural assays. The 7 strains used in this study were: 12 C57BL/6N (CR #027), 8 C57BL/6J (in house colony JAX #000664), 7 BALB/cJ (JAX #000651), 8 DBA/2J (JAX #000671), 8 BTBR, 8 FVB/NJ (JAX #001800), 8 A/J (JAX #000646). There were fifty-nine 10-12 weeks old male adult mice in total. All mice were injected with MnCl2 contrast agent and underwent whole brain magnetic resonance imaging two days before the start of the home-cage recording. Following two weeks of home-cage recording, all mice underwent open field testing followed by the three-chamber task two days later.

A second, independent cohort consisting of only C57BL/6J and BTBR strains was used to characterize home-cage social behaviour under conditions that were not influenced by prior imaging or behavioural testing. These strains were selected because they exhibit robust differences in sociability, and the sample size for each strain was increased to improve statistical power for analyses of home-cage metrics. Cohort 2 consisted of 6-week-old adult males that were housed in groups of 3 or 4 per cage. C57BL/6J (JAX #000664) were obtained from an in-house colony and BTBR mice (JAX #002282) were delivered from Jackson labs at three weeks of age to allow for time to acclimate to the housing rooms. A total of 42 mice (28 C57BL/6J and 14 BTBR) were used.

All analyses validating home-cage metrics and circadian dynamic analysis were performed using cohort 2, whereas analyses examining correlations of home-cage metrics with neuroanatomical measures, three-chamber social behaviour, and open field assessment were conducted using cohort 1.

### Surgical implantation

All mice were injected with radiofrequency identification device (RFID) microchips the day before the first night of recording. RFID chips (Biomark BioTherm 13) were purchased from Merck Animal Health (https://www.merck-animal-health-usa.com) and implanted subcutaneously in the abdomen area of the animal to optimize detection from the antenna baseplate below the cage. Mice were anesthetized with isoflurane, and chips were implanted on the right side of the sagittal midplane following sanitization of the implantation site with 70% isopropyl alcohol. After implantation, but before recovery, an RFID reader was used to verify the chip was working and matched the ID number on the manufacturer packaging. The injection site was then stitched using absorbable stitches to prevent the chips from falling out. After recovery, mice were returned to their home cage and cages were placed on the HCM rack for tracking. Any chips that malfunctioned or fell out during the study would be replaced with a new chip implanted on the left side using the same procedure.

### Manganese-enhanced Magnetic Resonance Imaging (MEMRI) and image registration

One day before the mice in the first cohort were implanted with RFID chips, they underwent *in vivo* MR imaging. Twenty-four hours prior to MR scans, mice were injected intraperitoneally with manganese chloride contrast agent (0.04mmol/kg dose of 30 mM MnCl_2_ solution). Neuroanatomical images were acquired using a multi-channel 7.0 Tesla MRI scanner with a 30 cm diameter bore (Bruker, Billerica, Massachusetts), which enabled simultaneous imaging of up to four mice using cryogenically cooled surface coils.^48^ During the scan, mice were kept warm using a heated water platform and were anesthetized with 1-2% isoflurane in 100% oxygen. Respirations were continuously monitored using self-gated signals.^49^ Images were acquired using a three-dimensional T1-weighted FLASH gradient-echo sequence with the following parameters: TR = 26 ms, TE = 8.25 ms, flip angle = 23°, field of view = 25 × 22 × 22 mm, matrix size = 334 × 294 × 294, and two averages, resulting in an isotropic spatial resolution of 75 μm. Following acquisition, MR images were corrected for gradient field inhomogeneity-induced distortions and intensity non-uniformity.^50,51^

Images were registered using an automatic registration pipeline within the PydPiper toolkit to align images to a consensus average space.^52^ Individual images were quality checked for correct orientation and adequate contrast and resolution prior to running MAGeT. Structural brain volume changes between mice were determined using the MAGeT Brain algorithm,^53^ with a pre-segmented input atlas containing 159 brain structures.^54–56^ MAGeT outputs were quality controlled by overlying the atlas on top of brain images and ensuring the correct segmented regions are mapped onto each brain structure.

### Home-Cage monitoring (HCM) and information theory computation

Mouse social behaviour was recorded in Actual Analytics Home Cage Analyser (https://www.actualanalytics.com/). RFID-implanted mice were continuously tracked using 18 antennas arranged in a 6 x 3 grid beneath the cage floor, with approximately 50 mm between adjacent antenna positions. Mice were recorded for 14 nights, with one cage change performed during the day following the first week. Although data were collected during both light and dark phases, analyses were restricted to the dark phase to capture behaviour during the mice’s active period. Positional time series were subsequently processed to quantify spatial occupancy, movement and information-theoretic measures of social behaviour, as described below. Converting time series data of mouse positions into measures of social behaviour is not trivial. While pairwise separation distance may provide an intuitive measure of social behaviour based on proximity, it cannot distinguish social attraction or avoidance from coincidental co-occupancy or shared spatial preferences. For example, two mice may occupy the same region because one is attracted to the other, because both prefer the same location, or because of a shared response to an environmental stimulus. These limitations motivate an approach that quantifies the statistical dependence between the movements of cagemates rather than relying on proximity alone.

Information theory provides a framework for quantifying this dependence directly from positional trajectories. To quantify this dependence, we first characterize the uncertainty in each mouse’s positional distribution using Shannon entropy. Let X denote the position of a mouse, with possible positions 1 through 18, corresponding to the antenna locations along the home cage. The entropy of X, measured in nats, is given by:

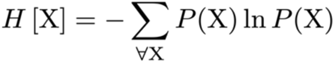

Since the uncertainty of a mouse’s location is measured by the entropy of the distribution of its locations, the dependency between mice can be calculated by the reduction of entropy (or increase in knowledge) by knowing the location of another mouse. The joint entropy of X and Y is defined as:

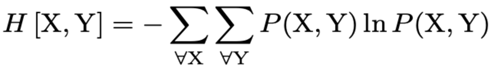

*H* [X,Y] represents the entropy (uncertainty) of the whole *X* and *Y* system, with *H* [X] and *H*[Y] representing the individual entropies corresponding to the marginal distributions of X and Y, respectively. If X and Y are independent, then the entropy of the whole system (joint entropy) would be the sum of the entropies of the individuals. On the other hand, if X and Y were dependent, the joint entropy would be less than the sum of individual entropies. It is this difference between the sum of the individual entropies and joint entropy that is known as mutual information (MI).

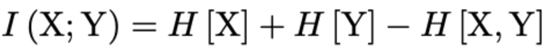

MI was computed in the natural unit of information (nats), with a value of 0 indicating the positions of two mice are independent. Higher MI values indicate greater shared information between the positions of the mice, such that knowing the position of one mouse provides more information about the position of the other. Thus, higher MI can be interpreted as greater statistical dependence between mice and, in the context of this study, greater social interaction. Since mice were housed in groups, social behavior was not limited to pairwise interactions. We therefore extended MI to quantify the information shared between the position of a mouse and the joint positions of its cagemates. For example, in the context of 3 mice in a cage, we can quantify the effect of Mouse B on Mouse A, but this time conditioning for the effect of Mouse C’s position. Thus, the MI between X and Y conditioned on Z is:

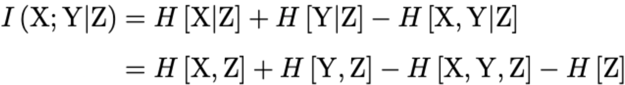

Rather than considering the effect of each variable independently, we used the concept of conditional MI to calculate the knowledge gained about Mouse A by knowledge of both Mouse B *and* Mouse C. For random variables X, Y, and Z, the knowledge gained about X knowing both Y and Z is:

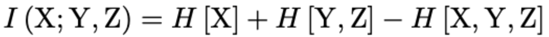

This concept can be extended to any number of variables. For instance, in a cage of n +1 mice, whose positions are given by random variables X_i_, where i goes from 0 to n. The information gained about the 0th mouse from knowing its cagemates is:

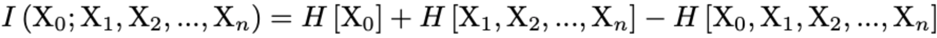

In this way, MI can be calculated between each mouse’s position and the joint positions of its cagemates, using the information gained about an individual’s location from the positions of the other mice as a measure of social behaviour. For each mouse, MI was computed at each prespecified temporal window (30, 60, 300, and 90 s) across the recording period. To obtain a cage-level estimate, MI observations were pooled across mice and recorded nighttime intervals and averaged within each cage for each temporal window. These cage-level estimates were then used in comparisons with null distributions.

To investigate the temporal nature of information transfer, we also computed transfer entropies which quantify how much information flows from X to other random variables (how much does a mouse’s location preference affect its cagemates’ location preferences), or from other variables into X (how much does the cagemates’ location preferences affect a mouse’s location preference). In other words, transfer entropy helps us determine whether social hierarchies exist in a cage by determining if mice influence the future behavior of their cagemates. Transfer entropy (TE) is defined as MI between variable X at time *t* and variable Y at an earlier time *t - L*, conditioned on X at an earlier time.

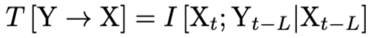

Information-theoretic measures can be influenced by temporal parameter selection, group size, boundary effects within the home cage, differences in free-running periods, and estimator bias. We therefore implemented a null-model framework to establish the expected baseline for MI and TE under randomized cage membership while preserving strain-specific movement dynamics. Null datasets were generated from strain-matched cage permutations by constructing artificial “null cages” composed of either three or four unrelated mice from different physical cages but of the same strain (100 null cages per strain of 50 cages per group size). Duplicate cage compositions and duplicate mice within a cage excluded. MI and TE were pooled and averaged within each real and null cage for each temporal window (30, 60, 300, and 900 s) and group size (three or four mice), such that each cage contributed one value per temporal window.

Analyses were restricted to C57BL/6J, which had 8 real cages: 4 three-mouse and 4 four-mouse cages, each analyzed at all 4 temporal windows. For each combination of temporal window and group size, the test statistic was the mean cage-level MI (or TE) across the 4 real cages. To test whether real cages differed from chance, we asked how often a result as extreme as ours would occur if cage membership did not matter. The null distribution was built from the strain-matched null cages with the same number of mice per cage. We computed the mean cage-level MI (or TE) of every possible combination of 4 null cages (C(50, 4) = 230,300 combinations), which gives the full range of means expected by chance (exact resampling test against a permutation-derived null). We then measured how far the real mean was from the average of all null cages (the real gap), how far each of the 230,300 null means was from that same average (the null gaps), and counted how many null gaps were at least as large as the real gap, in either direction. The two-sided p-value was computed as (1 + number of null gaps ≥ real gap) / (230,300 + 1). The +1 treats the observed result as one of the possible outcomes under the null, so p cannot equal zero. P-values were adjusted across the 8 comparisons for MI and TE (4 windows × 2 group sizes) using the Benjamini-Hochberg procedure. As a standardized effect size, we also report Z = (mean of real cages - mean of all null cages) / standard deviation of cage-level values across null cages. Z was not used for inference. Analyses were restricted to C57BL/6J. BTBR cages were collected but excluded from this analysis because only two real cages per group size were available. This approach allowed us to determine whether observed information-theoretic measures exceeded those expected from chance co-occurrence and strain-specific movement patterns alone, thereby providing evidence that the measured interactions reflected genuine social behaviour.

We also evaluated the influence of temporal parameters on information theory metrics by systemically varying the bin time (30 seconds (s), 60s, 300s, 900s) and lag time parameters (for TE) to compare the resulting probability distribution between our real and null data. As movement patterns in individual mice can vary substantially across timescales, ranging from brief bursts of activity over seconds to slower behavioral changes occurring over minutes or hours, we compared our real and null distribution averaged by mouse and by cage. These analyses were used to identify the optimal temporal parameters that maximized separation between real and null distributions for downstream analysis. All analyses were performed using a 300 second (5-minute) time bins.

To assess the sensitivity and robustness of our MI estimates, we additionally computed MI using the shrinkage entropy estimator implemented in the *infotheo* R package (method = “shrink”). Unlike the default estimator used in our primary analysis (method = “emp”), which calculates probabilities directly from observed frequencies, the shrinkage estimator uses a regularized estimate of the underlying probability distribution. It achieves this by shrinking the observed probabilities toward a uniform distribution using a shrinkage weighting term (λ). This provides an alternative approach to estimating entropy when the number of possible joint states is large relative to the number of observations. For each focal mouse, MI was calculated using the same formulation as the actual estimator, but with shrinkage-based entropy estimates for the focal mouse, the joint state of the remaining mice, and the joint state of all mice. The shrinkage estimator was applied consistently to both real and null data, allowing us to assess whether the observed differences between these datasets were sensitive to the choice of entropy estimator.

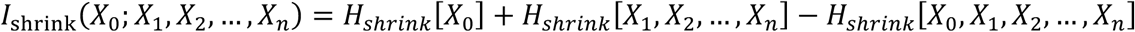

Unless otherwise stated, metrics were analyzed using a linear mixed-effects model implemented in the lmerTest 3.1.3 package in R statistical software (version 3.6.3). Given the longitudinal time series nature of this dataset, we accounted for repeated measures within the same subject by including a random intercept of pairwise interaction. Significance testing was conducted using t-tests with Satterthwaite’s method. Post hoc comparisons for the effect of number of mice in a cage (cage size) were conducted using estimated marginal means via the *emmeans* R package, with pairwise comparisons computed for cage size. To estimate uncertainty in all observed metrics, we applied a non-parametric bootstrap procedure with 1000 resamples of the data. For each resample, entropy and MI were recalculated, and 95% confidence intervals were derived from the bootstrap distribution.

To validate MI against direct observation, manually scored social behaviour counts from video clips (n = 24 5-minute clips from 6 representative cages) were compared with MI using Pearson correlation within each strain. Since counts are non-negative and over dispersed, we additionally modeled our data with a negative binomial generalized linear mixed model (log link) fit by maximum likelihood (Laplace approximation) using the glmer.nb function in the *lme4* package (version 1.1.34) in R. The model included MI and strain (C57BL/6J as reference) as fixed effects and a random intercept for cage to account for multiple clips from the same cage. We tested whether the MI and behaviour counts relationship differed between strains using a likelihood ratio test comparing models with and without an MI × strain interaction. Effects are reported as rate ratios (exponentiated coefficients) with Wald 95% confidence intervals.

### Behavioural Testing

#### Three-chamber test

The three-chamber test experiment design consisted of a habituation and trial phase. During the habituation phase, the test mouse was placed in the center chamber and could freely explore the other two adjacent chambers. After 10 minutes of habituation, the test mouse was placed in the center chamber. A novel mouse was introduced to the one chamber under a wired cup while an identical cup in the adjacent chamber remained empty. The mouse was then given access to the adjacent chambers and time spent in each chamber was assessed. Sociability index (SI) is defined as the ratio of the difference of time spent in the chamber with the novel mouse and the chamber with the empty cup over the sum of time in both chambers. In this study, age- and sex-matched 129S1/SvImJ (JAX #002448) mice were used as the novel stimulus because of their relatively low activity levels, minimal aggression, and docile temperament. Stranger mice were used with a maximum of two test mice and were not used in adjacent trials. After each trial, both the test and stranger mice were returned to their respective cages, feces and urine were removed, and the entire apparatus was disinfected with Clidox before beginning the next trial.

#### Open field test

The open field test (OFT) was used in this study as a measure of anxiety, locomotor and exploratory behaviours. Animals were transported to the testing room and left undisturbed for 30 minutes before the test. Mice were individually placed in the center of an open-field arena (SuperFlex 42×42×30cm under 200 LUX) and allowed to freely explore the apparatus for the duration of the trial (20 minutes). Animal movement was recorded using an overhead camera and analyzed using automated tracking software (Fusion software by Omnitech Electronics, Inc).

To quantify the spatial distribution of open-field exploration (entropy), the arena was divided into sixteen 4” x 4” zones using a custom zone map. The duration spent in each zone was obtained for consecutive 5-min intervals throughout the 20-min test. Within each interval, zone occupancy probabilities were calculated as the proportion of the total recorded time spent in each of the 16 zones. Shannon entropy was calculated from these occupancy probabilities (same as above), providing a measure of how evenly the mouse distributed its time across the available spatial states. Entropy was normalized by the maximum entropy for a 16-zone environment, yielding values ranging from 0 to 1, with higher values indicating a more even distribution of time across spatial zones.

## Supporting information

Supplementary Video 1: High MI C57BL6J

Supplementary Video 2: Low MI C57BL6J

Supplementary Video 3: High MI BTBR

Supplementary Video 4: Low MI BTBR

## REFERENCES

1. Baribeau, D. A., Dupuis, A., Paton, T. A., Hammill, C. & Scherer, S. W. Structural neuroimaging correlates of social deficits are similar in autism spectrum disorder and attention-deficit/hyperactivity disorder: analysis from the POND Network. Transl Psychiatry 9, 72 (2019).

2. Dodell-Feder, D., Tully, L. M. & Hooker, C. I. Social impairment in schizophrenia: new approaches for treating a persistent problem. Curr Opin Psychiatry 28, 236–242 (2015).

3. Choi, T. Y., Jeong, S. & Koo, J. W. Mesocorticolimbic circuit mechanisms of social dominance behavior. Exp Mol Med 56, 1889–1899 (2024).

4. Jabarin, R., Netser, S. & Wagner, S. Beyond the three-chamber test: toward a multimodal and objective assessment of social behavior in rodents. Mol. Autism 13, 41 (2022).

5. Moy, S. S., Nadler, J. J., Young, N. B., Nonneman, R. J. & Grossman, A. W. Social approach in genetically engineered mouse lines relevant to autism. Genes Brain Behav 8, 129–142 (2009).

6. Nadler, J. J., Moy, S. S., Dold, G., Simmons, N. & Perez, A. Automated apparatus for quantitation of social approach behaviors in mice. Genes Brain Behav 3, 303–314 (2004).

7. Yang, M., Silverman, J. L. & Crawley, J. N. Automated three-chambered social approach task for mice. Curr Protoc Neurosci 56, 8 26 1–8 26 16 (2011).

8. Crawley, J. N. Designing mouse behavioral tasks relevant to autistic-like behaviors. Ment Retard Dev Disabil Res Rev 10, 248–258 (2004).

9. Shemesh, Y., Benjamin, A., Shoshani-Haye, K., Yizhar, O. & Chen, A. Studying dominance and aggression requires ethologically relevant paradigms. Curr Opin Neurobiol 86, 102879 (2024).

10. Sofyana, M., Piggins, H. D., Jackson, M. G. & Robinson, E. S. J. Beyond the barrier: an assessment of social behaviour in mice using a modified 3-chamber test. Physiol Behav 315, 115381 (2026).

11. Silverman, J. L., Yang, M., Lord, C. & Crawley, J. N. Behavioural phenotyping assays for mouse models of autism. Nat. Rev. Neurosci. 11, 490–502 (2010).

12. Szabó, J., Renczés, E., Borbélyová, V., Ostatníková, D. & Celec, P. Assessing sociability using the Three-Chamber Social Interaction Test and the Reciprocal Interaction Test in a genetic mouse model of ASD. Behav Brain Funct 20, 24 (2024).

13. Dhamne, S. C., Silverman, J. L., Super, C. E., Lammers, S. H. T. & Hameed, M. Q. Replicable in vivo physiological and behavioral phenotypes of the Shank3B null mutant mouse model of autism. Mol. Autism 8, 26 (2017).

14. Ey, E., Yang, M., Katz, A. M., Woldeyohannes, L. & Silverman, J. L. Absence of deficits in social behaviors and ultrasonic vocalizations in later generations of mice lacking neuroligin4. Genes Brain Behav. 11, 928–941 (2012).

15. Weissbrod, A., Shapiro, A., Vasserman, G., Edry, L. & Dayan, M. Automated long-term tracking and social behavioural phenotyping of animal colonies within a semi-natural environment. Nat. Commun. 4, 2018 (2013).

16. Iannello, F. Non-intrusive high throughput automated data collection from the home cage. Heliyon 5,**4**, (2019).

17. Chaumont, F., Coura, R. D., Serreau, P., Cressant, A. & Chabout, J. Computerized video analysis of social interactions in mice. Nat Methods 9, 410–417 (2012).

18. Mingrone, A., Kaffman, A. & Kaffman, A. The promise of automated home-cage monitoring in improving translational utility of psychiatric research in rodents. Front Neurosci 14, 618593 (2020).

19. Jhuang, H., Garrote, E., Mutch, J., Yu, X. & Khilnani, V. Automated home-cage behavioural phenotyping of mice. Nat Commun 1, 68 (2010).

20. Krackow, S., Vannoni, E., Codita, A., Mohammed, A. H. & Cirulli, F. Consistent behavioral phenotype differences between inbred mouse strains in the IntelliCage. Genes Brain Behav 9, 722–731 (2010).

21. Robinson, L., Spruijt, B. & Riedel, G. Between and within laboratory reliability of mouse behaviour recorded in home-cage and open-field. J Neurosci Methods 300, 10–19 (2018).

22. Crawley, J. N. Mouse behavioral assays relevant to the symptoms of autism. Brain Pathol 17, 448–459 (2007).

23. Moretti, P., Bouwknecht, J. A., Teague, R., Paylor, R. & Zoghbi, H. Y. Abnormalities of social interactions and home-cage behavior in a mouse model of Rett syndrome. Hum Mol Genet 14, 205–220 (2005).

24. Ohayon, S., Avni, O., Taylor, A. L., Perona, P. & Egnor, S. E. R. Automated multi-day tracking of marked mice for the analysis of social behaviour. J Neurosci Methods 219, 10–19 (2013).

25. Shannon, C. E. A mathematical theory of communication. Bell Syst Tech J 27, 379–423, 623– 656 (1948).

26. Shemesh, Y., Sztainberg, Y., Forkosh, O., Shlapobersky, T. & Chen, A. High-order social interactions in groups of mice. eLife 2, 00759 (2013).

27. Yang, M., Abrams, D. N., Zhang, J. Y., Weber, M. D. & Katz, A. M. Low sociability in BTBR T+tf/J mice is independent of partner strain. Physiol Behav 107, 649–662 (2012).

28. Ko, J. Neuroanatomical Substrates of Rodent Social Behavior: The Medial Prefrontal Cortex and Its Projection Patterns. Front. Neural Circuits 11, 41 (2017).

29. Li, H., Zhao, Z., Jiang, S. & Wu, H. Brain circuits that regulate social behavior. Mol. Psychiatry 30, 3240–3256 (2025).

30. Griguoli, M. & Pimpinella, D. Medial septum: relevance for social memory. Front Neural Circuits 16, 965172 (2022).

31. Kahnau, P., Mieske, P., Wilzopolski, J., Kalliokoski, O. & Mandillo, S. A systematic review of the development and application of home cage monitoring in laboratory mice and rats. BMC Biol 21, 256 (2023).

32. Grieco, F., Bernstein, B. J., Biemans, B., Bikovski, L. & Burnett, C. J. Measuring behavior in the home cage: study design, applications, challenges, and perspectives. Front Behav Neurosci 15, 735387 (2021).

33. Schreiber, T. Measuring information transfer. Phys Rev Lett 85, 461–464 (2000).

34. Timme, N. M. & Lapish, C. A tutorial for information theory in neuroscience. eNeuro 5, 0052–18 2018 (2018).

35. Scattoni, M. L., Martire, A., Cartocci, G., Ferrante, A. & Ricceri, L. Reduced Social Interaction, Behavioural Flexibility and BDNF Signalling in the BTBR T+ Tf/J Strain, a Mouse Model of Autism. (Behav Brain Res, 2013).

36. Fairless, A. H., Shah, R. Y., Guthrie, A. J., Li, H. & Brodkin, E. S. Deconstructing sociability, an autism-relevant phenotype, in mouse models. Anat. Rec. 294, 1713–1725 (2011).

37. Fairless, A. H., Katz, J. M., Vijayvargiya, N., Dow, H. C. & Kreibich, A. S. Development of home cage social behaviors in BALB/cJ vs. C57BL6J Mice Behav Brain Res 237, 338–347 (2012).

38. Mei, L., Osakada, T. & Lin, D. Hypothalamic control of innate social behaviors. Science 382, 399–404 (2023).

39. Menon, R., Süß, T., Oliveira, V. E. M., Neumann, I. D. & Bludau, A. Neurobiology of the lateral septum: regulation of social behavior. Trends Neurosci 45, 27–40 (2022).

40. Thompson, S. M., Berkowitz, L. E. & Clark, B. J. Behavioral and neural subsystems of rodent exploration. Learn Motiv 61, 3–15 (2018).

41. Heiney, S. A., Kim, D., Augustine, G. J. & Medina, J. F. Kinematics and dynamics of the mouse cerebellum during locomotion. Neuron 84, 1127–1137 (2014).

42. Shi, H.-J., Wang, S., Wang, X.-P., Zhang, R.-X. & Zhu, L.-J. Hippocampus: molecular, cellular, and circuit features in anxiety. Neurosci Bull 39, 1009–1026 (2023).

43. Arakawa, H., Blanchard, D. C. & Blanchard, R. J. Colony formation of C57BL/6J mice in visible burrow system: Identification of eusocial behaviors in a background strain for genetic animal models of autism. Behav Brain Res 176, 27–39 (2007).

44. Vijaya Shankara, J., Horsley, K. G., Cheng, N., Rho, J. M. & Antle, M. C. Circadian responses to light in the BTBR mouse. J Biol Rhythms 37, 498–515 (2022).

45. Terranova, M. L., Laviola, G. & Alleva, E. Ontogeny of amicable social behavior in the mouse: gender differences and ongoing isolation outcomes. Dev Psychobiol 26, 467–481 (1993).

46. Aubry, A. V., Burnett, C. J., Goodwin, N. L., Li, L. & Navarrete, J. Sex differences in appetitive and reactive aggression. Neuropsychopharmacology 47, 1746–1754 (2022).

47. Melotti, L., Kästner, N., Eick, A. K., Schnelle, A. L. & Palme, R. Can live with ‘em, can live without ‘em: Pair housed male C57BL/6J mice show low aggression and increasing sociopositive interactions with age, but can adapt to single housing if separated. Appl Anim Behav Sci 214, 79–88 (2019).

48. Arbabi, A., Spencer Noakes, L., Vousden, D., Dazai, J. & Spring, S. Multiple-mouse magnetic resonance imaging with cryogenic radiofrequency probes for evaluation of brain development. Neuroimage 252, 119008 (2022).

49. Nieman, B. J., Szulc, K. U. & Turnbull, D. H. Three-dimensional, in vivo MRI with self-gating and image coregistration in the mouse. Magn Reson Med 61, 1148–1157 (2009).

50. Nieman, B. J. et al. MRI to assess neurological function. Curr Protoc Mouse Biol 8, 44 (2018).

51. Sled, J. G., Zijdenbos, A. P. & Evans, A. C. A nonparametric method for automatic correction of intensity nonuniformity in MRI data. IEEE Trans Med Imaging 17, 87–97 (1998).

52. Friedel, M., Eede, M. C., Pipitone, J., Chakravarty, M. M. & Lerch, J. P. Pydpiper: a flexible toolkit for constructing novel registration pipelines. Front Neuroinform 8, 67 (2014).

53. Chakravarty, M. M., Steadman, P., Eede, M. C., Calcott, R. D. & Gu, V. Performing label-fusion-based segmentation using multiple automatically generated templates. Hum Brain Mapp 34, 2635–2654 (2013).

54. Dorr, A. E., Lerch, J. P., Spring, S., Kabani, N. & Henkelman, R. M. High resolution three-dimensional brain atlas using an average magnetic resonance image of 40 adult C57Bl/6J mice. Neuroimage 42, 60–69 (2008).

55. Steadman, P. E., Ellegood, J., Szulc, K. U., Turnbull, D. H. & Joyner, A. L. Genetic effects on cerebellar structure across mouse models of autism using a magnetic resonance imaging atlas. Autism Res 7, 124–137 (2014).

56. Ullmann, J. F. P., Watson, C., Janke, A. L., Kurniawan, N. D. & Reutens, D. C. A segmentation protocol and MRI atlas of the C57BL/6J mouse neocortex. Neuroimage 78, 196–203 (2013).

