## Supplementary Material for "An information theoretic approach to evaluating social behaviour in mice using automated home-cage monitoring"

A

| Window (s) | Mice per cage | Real cages | Null cages | Real mean MI | Null mean MI | Z | Permutation p | BH-adj. p |
|---|---|---|---|---|---|---|---|---|
| 30 | 4 | 4 | 50 | 0.193 | 0.154 | 2.34 | <0.0001 | <0.0001 |
| 30 | 3 | 4 | 50 | 0.169 | 0.112 | 5.10 | <0.0001 | <0.0001 |
| 60 | 4 | 4 | 50 | 0.341 | 0.278 | 2.42 | <0.0001 | <0.0001 |
| 60 | 3 | 4 | 50 | 0.291 | 0.208 | 4.51 | <0.0001 | <0.0001 |
| 300 | 4 | 4 | 50 | 0.871 | 0.734 | 2.73 | <0.0001 | <0.0001 |
| 300 | 3 | 4 | 50 | 0.697 | 0.570 | 2.94 | <0.0001 | <0.0001 |
| 900 | 4 | 4 | 50 | 1.29 | 1.10 | 2.96 | <0.0001 | <0.0001 |
| 900 | 3 | 4 | 50 | 0.972 | 0.850 | 1.95 | <0.0001 | <0.0001 |

C57BL/6J only. P-values are from an exact resampling test against the strain-matched null (two-sided), BH-adjusted across the 8 comparisons.
Z is a standardized effect size and was not used for inference.

B

| Window (s) | Mice per cage | Real cages | Null cages | Real mean TE | Null mean TE | Z | Permutation p | BH-adj. p |
|---|---|---|---|---|---|---|---|---|
| 30 | 4 | 4 | 50 | 0.0805 | 0.0746 | 0.91 | 0.054 | 0.087 |
| 30 | 3 | 4 | 50 | 0.0757 | 0.0609 | 2.79 | <0.0001 | **<0.0001** |
| 60 | 4 | 4 | 50 | 0.125 | 0.121 | 0.46 | 0.351 | 0.468 |
| 60 | 3 | 4 | 50 | 0.122 | 0.106 | 1.90 | 0.0001 | **0.0003** |
| 300 | 4 | 4 | 50 | 0.258 | 0.273 | −1.08 | 0.022 | **0.044** |
| 300 | 3 | 4 | 50 | 0.286 | 0.282 | 0.23 | 0.634 | 0.674 |
| 900 | 4 | 4 | 50 | 0.362 | 0.399 | −2.37 | <0.0001 | **<0.0001** |
| 900 | 3 | 4 | 50 | 0.471 | 0.467 | 0.20 | 0.674 | 0.674 |

C57BL/6J only. P-values are from an exact resampling test against the strain-matched null (two-sided), BH-adjusted across the 8 comparisons.
Z is a standardized effect size and was not used for inference. Bold: BH-adj. $p < 0.05$.

**Supplementary Table 1:** Comparison of real and null cage-level distributions A) mutual information (MI) and B) transfer entropy (TE) in C57BL/6J mice across temporal windows (30, 60, 300, and 900 s) and cage sizes (three- and four-mouse cages). For each cage, MI or TE was first averaged across nights, giving one value per cage. For each combination of temporal window and cage size, the mean of the 4 real cages was compared with a strain-matched null built from artificial cages of unrelated mice of the same strain (50 null cages per combination). Two-sided p-values were obtained from an exact resampling test: the real mean was compared with the mean of every possible set of 4 null cages (230,300 combinations), with a +1 correction, and adjusted across the 8 comparisons per measure using the Benjamini-Hochberg procedure (BH). Bold indicates BH-adj. $p < 0.05$. Z = (real mean − null mean) / null SD, where null mean and null SD are the mean and standard deviation of cage-level values across all null cages, is reported as a standardized effect size and was not used for inference.

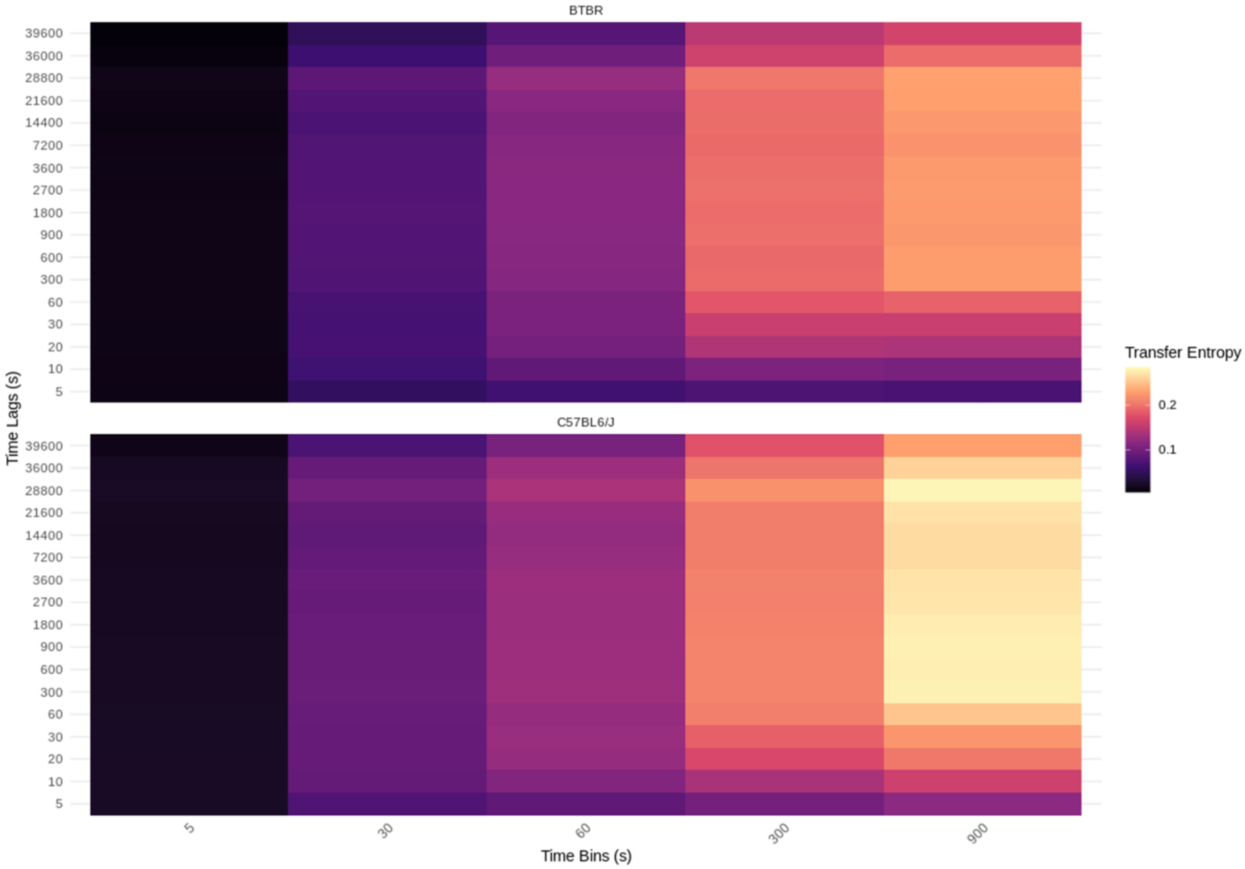


**Supplementary Figure 1:** Heatmap showing the range of transfer entropy values at varying time bins (in seconds) and time lags (in seconds). Time bin refers to the sampling time in which the data was averaged over while time lag refers to the delay in which one mouse's past position influences another. At 5 seconds sampling time, TE values were zero or near-zero and differences across lag times were negligible. An increase in TE values was observed with increasing time bins in both strains.

A

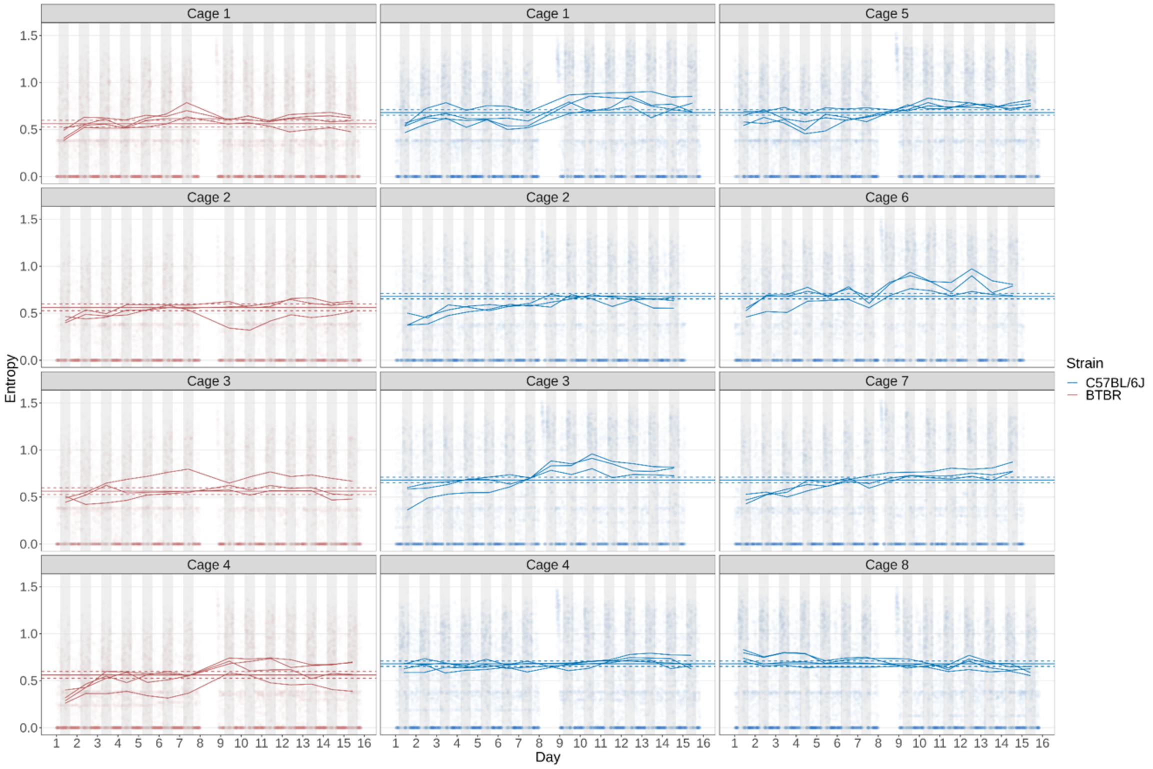


B

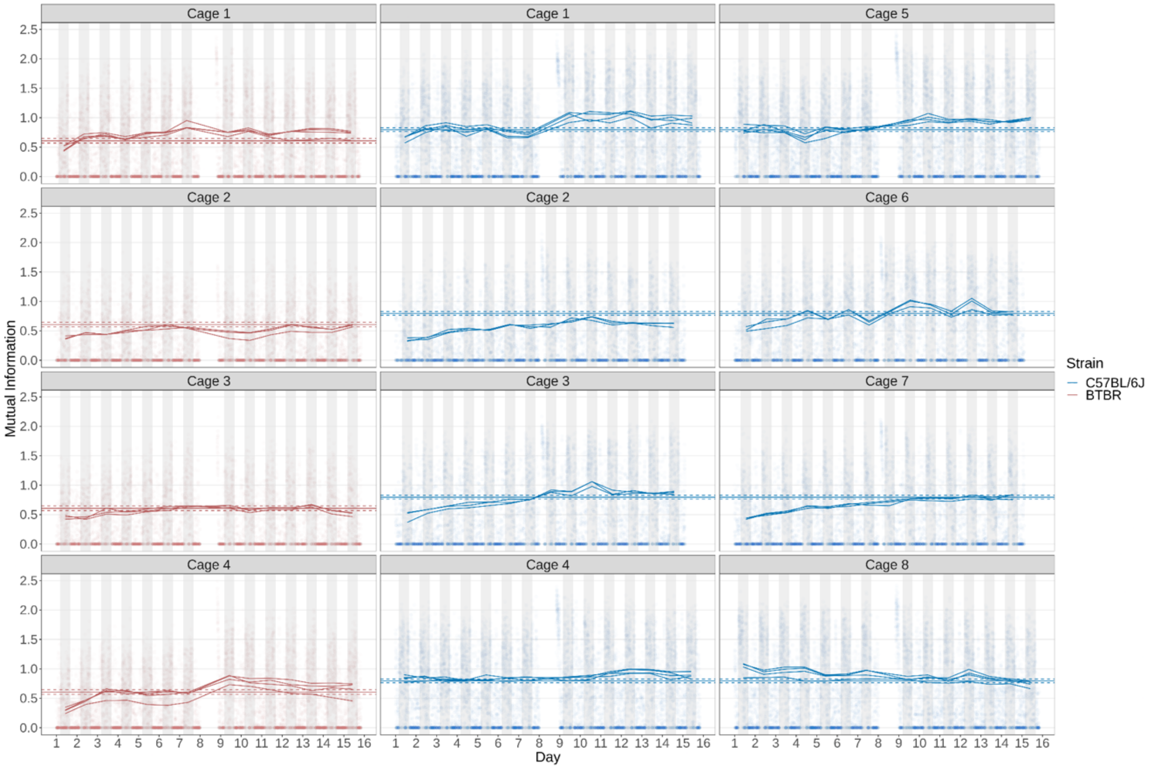


**Supplementary Figure 2: A)** Time series showing the full recording period (two weeks) for twelve cages, with the top and bottom rows each containing four mice, and the middle two rows containing three mice. **A)** Entropy per mouse was computed and averaged in aggregates of 5-minute time-bins (points). Data collected for each night (dark bars in the background) was averaged for each mouse and is given by the coloured lines. The horizontal lines represent the mean mutual information for each strain with the dashed lines representing 95% confidence intervals in the 1000-samples parametric bootstrap. **B)** Time series showing two weeks of mutual information (MI) recorded for twelve cages. Each point represents the MI between a mouse's position distribution and the joint position distribution of its cagemates averaged in aggregates of 5-minute time-bins.

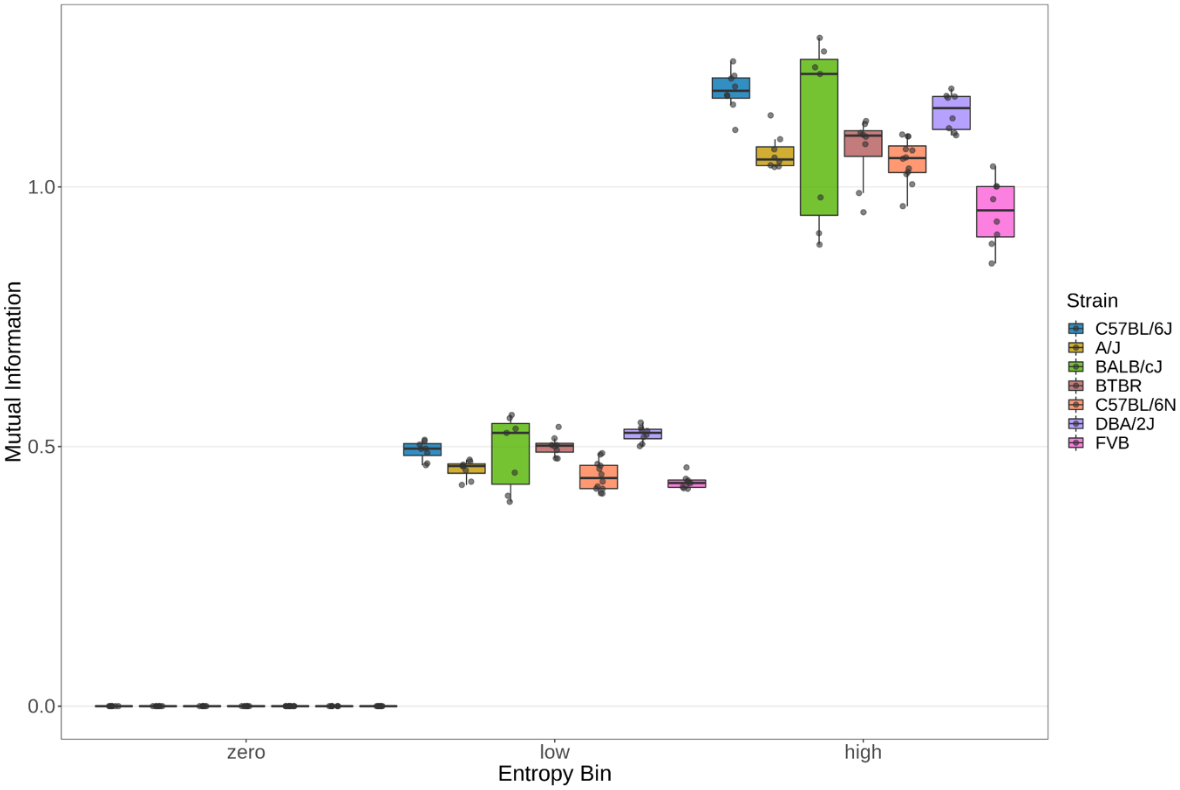


**Supplementary Figure 3:** Comparison of MI across strains at zero, low, and high entropy bins. Points represent average MI per mouse. Relative to our reference strain C57BL/6J, all other strains had a significantly lower increase in MI at the same increase in entropy.

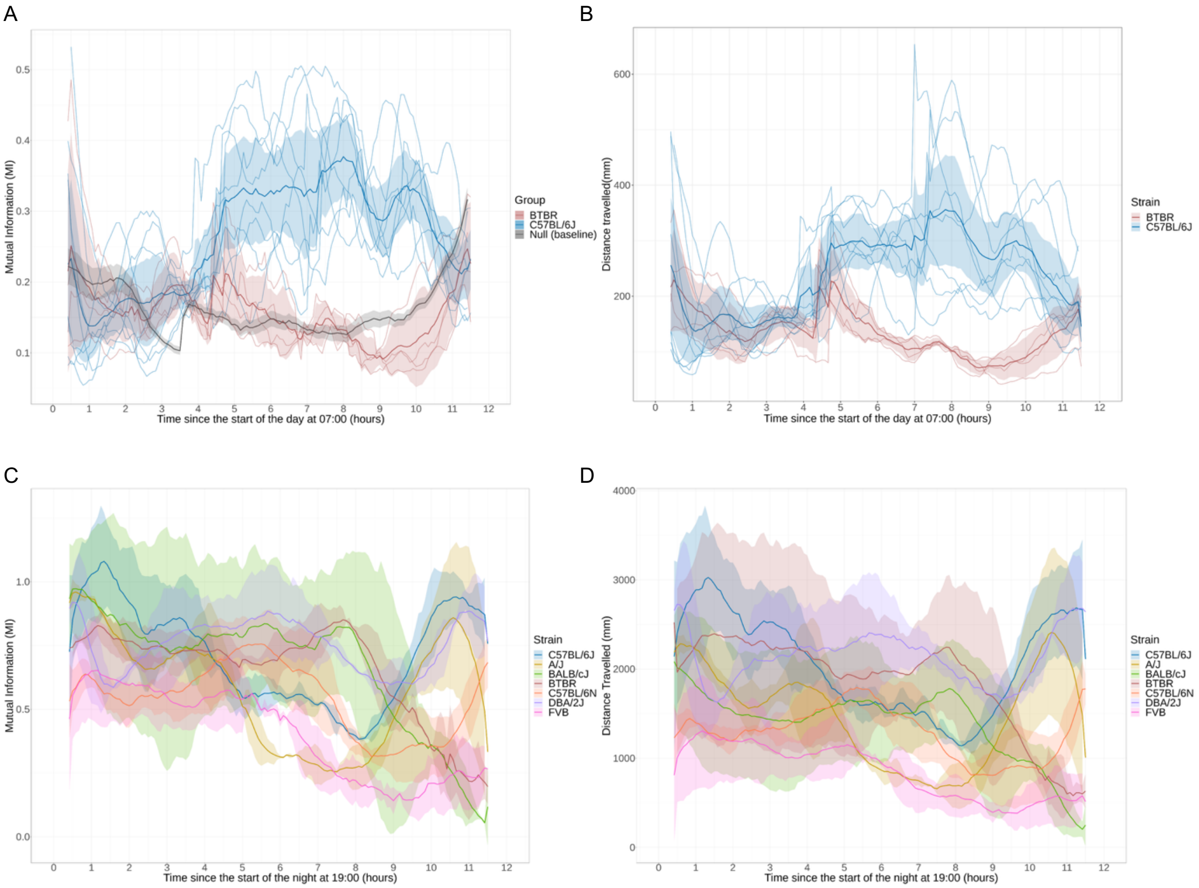


**Supplementary Figure 4:** Mouse activity during the inactive day cycle across different strains of mice. **A)** Sliding window estimates of MI between C57BL6/J and BTBR mice are plotted as a function of time since the start of the light phase (07:00) over a 12-hour period. Strain differences were observed in the second half of the day cycle with C57BL/6J showing elevated MI and Distance travelled than BTBR mice. For each 5-minute sampling time, MI was averaged over an hourly window centered on that timepoint, consisting of six preceding and six subsequent 5-minute bins each hour. Lines represent mean MI for each group (C57BL/6J, blue; BTBR, red; Null, grey), with shaded regions indicating 95% confidence interval. Thin lines show individual cage-level trajectories. **B)** Sliding window estimates of distance travelled averaged across one day cycle computed the same as MI. **C)** Sliding window estimates of MI between seven inbred strains. **D)** Sliding window estimates of distance travelled between seven inbred strains.

**Supplementary Table 2:** Correlation between brain structure volumes and all behavioural measures across mice. The table reports the t-statistic, correlation coefficient (Pearson's r), P value (p), and false discovery rate (FDR)-adjusted P value (q) for each ROI–behaviour association. FDR correction was performed across all ROI × behaviour comparisons (BH 10%).

**Supplementary Videos:** Representative C57BL/6J and BTBR cages with low and high MI corresponding to low and high frequency of observed social interactions respectively.
